# Disruption of basal CXCR4 oligomers impairs oncogenic properties in lymphoid neoplasms

**DOI:** 10.1101/2024.12.06.627249

**Authors:** Simon Mobach, Nick D. Bergkamp, Ziliang Ma, Marco V. Haselager, Stephanie M. Anbuhl, Daphne Jurriens, Jelle van den Bor, Ziming Wang, Caitrin Crudden, Jamie L. Roos, Claudia V. Perez Almeria, Rick A. Boergonje, Martin J. Lohse, Reggie Bosma, Eric Eldering, Marco Siderius, Wei Wu, Marcel Spaargaren, Sanne H. Tonino, Arnon P. Kater, Martine J. Smit, Raimond Heukers

## Abstract

The chemokine receptor CXCR4 is overexpressed in many cancers and contributes to pathogenesis, disease progression, and resistance to therapies. CXCR4 is known to form oligomers, but the potential functional relevance in malignancies remain elusive. Using a newly established nanobody-based BRET method, we demonstrate that oligomerization of endogenous CXCR4 on lymphoid cancer cell lines correlates with enhanced expression levels. Specific disruption of CXCR4 oligomers reduced basal cell migration and pro-survival signaling via changes in the phosphoproteome, indicating the existence of basal CXCR4-oligomer-mediated signaling. Oligomer disruption also inhibited growth of primary CLL 3D spheroids and sensitized primary malignant cells to clinically used Bcl-2 inhibitor venetoclax. Given its limited efficacy in some patients and the ability to develop resistance, sensitizing malignant B-cells to venetoclax is of clinical relevance. Taken together, we established a new, non-canonical and critical role for CXCR4 oligomers in lymphoid neoplasms and demonstrated that selective targeting thereof has clinical potential.

**Significance statement:** Class A GPCRs, including the chemokine receptor CXCR4, can form oligomers, but their functional relevance remains poorly understood. This study provides evidence for the role of basal CXCR4 oligomers in lymphoid neoplasms, where they drive pro-survival signaling, migration, and tumor growth. We use a novel nanobody-based BRET method to demonstrate that endogenous CXCR4 constitutively oligomerizes in lymphoid cancer cells, correlating with receptor expression levels. Pharmacological disruption of these oligomers reduces tumor-associated signaling, impairs spheroid growth, and sensitizes patient-derived malignant cells to the apoptosis-inducing drug Venetoclax. Since CXCR4 is frequently overexpressed and potentially clustered in various malignancies, this work offers broader implications for enhancing treatment efficacy, overcoming drug resistance, and potentially reducing side effects across multiple cancer types.

## Introduction

The treatment landscape of B cell malignancies like chronic lymphocytic leukemia (CLL) has undergone significant transformations after the introduction of effective oral targeted therapies such as BTK-, PI3K- and Bcl-2 inhibitors (Venetoclax) and next-generation anti-CD20 monoclonal antibodies (*1*). Nevertheless, the unresponsiveness of some patients, along with acquired resistance and the nearly universal subsequent relapse of the disease, underscores the ongoing need for a potentially curative treatment (*2*). The chemokine receptor CXCR4 is overexpressed in many human cancers, including lymphoid neoplasms. In these disease states, CXCR4 induces signaling that promotes tumor survival and metastasis upon activation by its endogenous ligand CXCL12 (*3*). In CLL, CXCR4 was shown to promote a protective tumor microenvironment by allowing migration into the lymph node and altering the behavior of adjacent cells to support tumor survival and growth (*4*). CXCR4 signaling also drives the retention of malignant cells in the bone marrow, thereby protecting these cells from chemotoxic stress, and targeted therapies administered (*5–7*). These observations support a central critical role for CXCR4 signaling in the biology of lymphoid neoplasms and positions CXCR4 as an important drug target to treat such diseases (*8, 9*).

CXCR4 belongs to the class A G protein-coupled receptors (GPCRs). Since GPCRs regulate numerous (patho-)physiological processes and are highly amenable to drug intervention, they represent a major class of drug targets (*10*). Although classically perceived as monomeric signaling units, GPCRs are increasingly recognized to exist and signal as dimers or as higher-order oligomeric complexes (*11*). For example, it is well-established that the functionality of class C GPCRs is critically dependent on the formation of homo- and heterodimers (*12, 13*). In contrast, the role of oligomerization in regulating receptor function and influencing downstream signaling outcomes remains unclear for the larger class A GPCR family. Some class A GPCRs appear to form transient dimers and higher-order oligomers (*14, 15*). However, the physiological roles of such complexes remain poorly understood to date.

A large body of evidence indicates that CXCR4 is capable of forming dimers and higher-order oligomers (i.e., clusters of three or more receptors) (*16–25*). Upon recombinant overexpression to levels mimicking an oncogenic setting, CXCR4 exists almost exclusively as dimers or higher-order oligomers (*18, 19*). In malignant lymphocytic T-cells, CXCR4 mainly resides in higher-order oligomers while it is largely monomeric in primary healthy T-cells (*24*), thus suggesting that CXCR4 oligomers exist and contribute to malignancy. Moreover, CXCL12-mediated migration of CXCR4-expressing T-cells was reported to be dependent on enhanced higher-order oligomer formation (*24, 26*), illustrating that CXCR4 oligomerization impacts receptor function. However, it remains to be clarified whether CXCR4 oligomerization drives malignancy. Additionally, it is important to investigate which specific pro-tumorigenic effects of CXCR4 are associated with signaling pathways unique to receptor oligomerization.

In this study, we set out to investigate the malignant potential of CXCR4 oligomerization. Using nanobody-based bioluminescence resonance energy transfer (BRET) and direct stochastic optical reconstruction microscopy (dSTORM) single-molecule imaging, we studied CXCR4 oligomerization in a panel of lymphoid neoplasm cell lines and primary cultures. Using mass spectrometry-based phosphoproteomics, we assessed signaling downstream of CXCR4 oligomers. Specific changes at the phosphosite level led us to uncover basal migration, spheroid growth and cell survival as phenotypic consequences of basal CXCR4 oligomer-mediated signaling. Moreover, the attenuation of these phenotypes obtained by pharmacologically disrupting CXCR4 oligomers suggests that such clusters can serve as a novel therapeutic target with clinical potential.

## Results

### Detection of endogenous CXCR4 oligomers using nanobody-based BRET approach

Oligomerization of CXCR4 has been extensively studied in heterologous expression systems (*18, 19*). To investigate whether this also occurs in a native setting, we developed a method for the detection of untagged GPCR oligomers in living cells. Such analysis requires a detection molecule that binds to CXCR4 with high affinity, without altering the basal oligomeric state of the receptor or its basal receptor signaling. One of the previously selected CXCR4-binding nanobodies (*27*), VUN415, displayed such properties (Table S1). To allow the detection of CXCR4 clusters by BRET, VUN415 was either genetically fused to NanoLuciferase (NanoLuc, Nluc) or conjugated to a fluorescent dye (Fig. 1A). Close proximity of two or more receptors enables BRET between nanobody donor and acceptor constructs bound to different CXCR4 protomers, thereby providing information about the relative receptor oligomeric state. Indeed, increasing equimolar concentrations of the two nanobody fusion constructs led to a concentration-dependent, saturable increase in nanobody binding on CXCR4-overexpressing HEK293T cells (Fig. S1A), as well as an increase in BRET ratio (Fig. 1A). This detection was CXCR4 specific, as no binding of the probes (Fig. S1A), and therefore no BRET, was observed in CHO-K1 or CRISPR Cas9 CXCR4-knockout HEK293T cells, which both lack CXCR4 expression (Fig. S1B). In addition, no BRET was observed in these CXCR4^negative^ models when using a fixed saturating nanobody concentration with varying donor:acceptor probe ratios (Fig. S1C, S1D). No BRET signals were observed when VUN415-ATTO565 was replaced by unlabeled VUN415 (Fig. S2). Collectively, these results indicate our BRET approach is highly specific, quantitative and can be used effectively to monitor the oligomerisation status of CXCR4 in subsequent mechanistic experiments.

**Figure 1.**
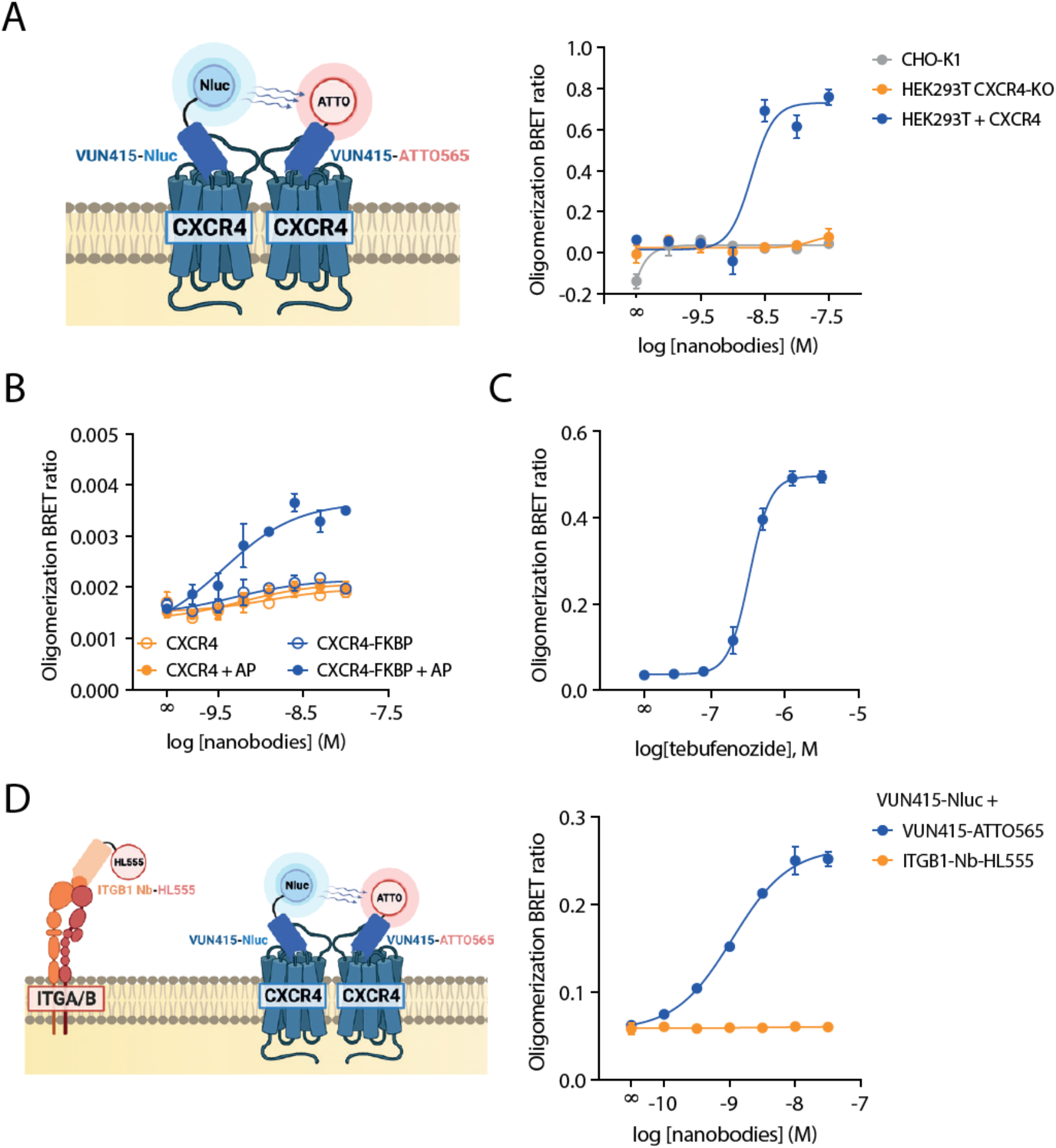
Detection of endogenous CXCR4 oligomers using nanobody-based BRET. **A** Schematics and results for the nanobody-based nanoBRET method to detect receptor oligomerization in CXCR4-overexpressing HEK293T, CHO-K1 and HEK293T CXCR4 CRISPR-Cas9 KO cells. Increasing equimolar concentrations of detection nanobodies VUN415-NanoLuc (‘Nluc’) and VUN415-ATTO565 were used. **B, C** Nanobody-based BRET measurement of receptor oligomerization using **(B)** untagged and FKBP-tagged CXCR4 or **(C)** ecdysone-inducible CXCR4. Stimulation with **(B)** 1 μM of dimerization ligand AP20187 (‘AP’) to induce dimerization or **(C)** increasing concentrations tebufenozide to induce receptor expression, as indicated. **D** Schematics and data of endogenous oligomer detection in Namalwa cells using VUN415-NanoLuc (‘Nluc’) as donor together with VUN415-ATTO565 or ITGB1-Nb-HL555 as acceptor. Data are mean ± SD and are representative of at least three independent experiments, each performed in triplicate.

Dimerization and higher-order oligomerization of proteins can be artificially induced through fusion to FK506-binding protein (FKBP) domains and subsequent chemical crosslinking (*28*). At low expression levels where CXCR4 is expected to be predominantly monomeric (*18–20*), a robust increase in nanobody oligomerization BRET was observed upon stimulation with crosslinker AP20187 for FKBP-tagged CXCR4 and not for the untagged receptor (Fig. 1B, S3). This shows that increased BRET values observed with our nanobody-based BRET approach is a consequence of receptor oligomerization. To verify that CXCR4 oligomerization depends on its expression level, as suggested previously (*18, 19*), an ecdysone-inducible CXCR4 expression construct was generated (*29*). Stimulation with ecdysone agonist tebufenozide indeed led to a concentration-dependent increase in CXCR4 expression, as well as nanobody-based oligomerization BRET signal (Fig. 1C and S4).

After the initial validation of the nanobody-BRET approach in HEK293T cells, we assessed the existence of endogenous CXCR4 oligomers. We focused on lymphoid neoplasms as enhanced CXCR4 levels are considered to play a prominent pathological role (*30–32*). Using the Namalwa Burkitt lymphoma cell line as a proof-of-concept, we observed a robust increase in BRET signal in cells treated with CXCR4 detection nanobodies, whereas no BRET occurred when combining VUN415-Nluc with a fluorescently labeled nanobody against the highly expressed integrin β1 (Fig. 1D). The lack of BRET for the integrin β1 control was not due to a lack of nanobody binding, as clear concentration-dependent binding to Namalwa cells was observed (Fig. S5). Hence, our nanobody-based BRET approach specifically showed the presence of heterologously and endogenously expressed CXCR4 oligomers.

### Enhanced oligomerization of endogenous CXCR4 on lymphoid cancer cell lines

Subsequently, we assessed whether CXCR4 oligomerization is associated with elevated expression of endogenous receptors on cancer cells. First, concentration-response curves for nanobody-based oligomerization detection were generated for a small selection of lymphoid cancer cell lines with varying CXCR4 expression levels and disease subtypes (Fig. 2A and S6). In the CLL cell line MEC-1 with very low CXCR4 expression, no oligomerization BRET signal was detected, confirming specificity of the BRET assay for cells other than the HEK293T CXCR4 CRISPR Cas9 KO and CHO-K1 cells. We observed concentration-dependent increases in BRET signal for CXCR4^low^ RPC1-WM1 (Waldenstrom macroglobulinemia) cells and CXCR4^high^ Z-138 (mantle cell lymphoma, MCL) cells, with similar BRET50 but much higher BRET_max_ values for the latter. Although different plate reader gain settings were used between cell lines, this did not affect the transformed BRET_max_ values (Fig. S7), and can therefore be considered to accurately reflect relative oligomerization levels. Out of a large and diverse panel of lymphoid neoplasms, oligomeric complexity of endogenous CXCR4 generally correlated to the receptor expression levels (Fig. 2B). Notably, some cell lines exhibited higher oligomeric complexity than expected based on their expression level, including cell lines PGA-1, L363 and CII. These deviations could be explained by other factors influencing oligomeric complexity, including membrane lipid composition (*33, 34*).

**Figure 2.**
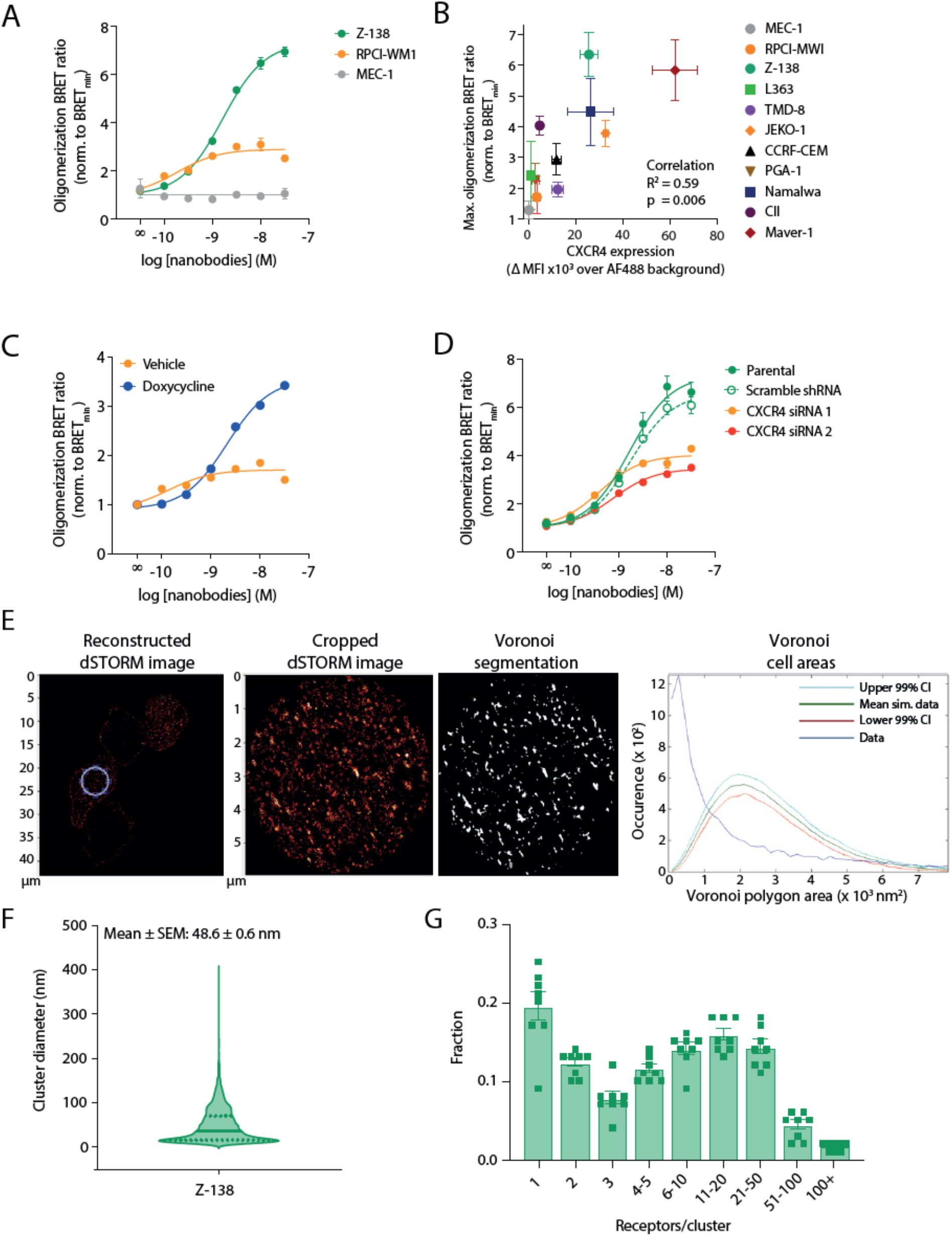
Differential oligomerization of endogenous CXCR4 receptors in lymphoid cancer cell lines. **A** Nanobody-based BRET measurement of CXCR4 oligomerization in lymphoid cancer cell lines MEC-1, RPCI-WM1 and Z-138. Data are representative of at least three independent experiments and depicted as mean ± SD. **B** Normalized CXCR4 oligomerization BRET_max_ plotted aginst flow cytometry surface receptor expression levels for lymphoid cancer cell line panel. Data are pooled mean ± SEM of at least three independent experiments, each performed in triplicate. **C, D** Effects on nanobody-based receptor oligomerization BRET of enhanced CXCR4 expression using doxycycline-inducible CXCR4 in RPCI-WM1 cells **(C)** or silencing of CXCR4 using scramble shRNA or CXCR4-targeting siRNA in Z-138 cells **(D)**. Data, normalized to BRET_min_ value of each individual cell line, are representative of at least three independent experiments and depicted as mean ± SD. **E** Data of dSTORM imaging and spatial point distribution analysis using Voronoi segmentation on Z-138 (CXCR4^high^) cells. Full reconstructed dSTORM image and analyzed region of image (‘Cropped dSTORM’), indicated by blue circle, are visualized. Corresponding thresholded binary map and Voronoi polygron area plot are shown. In Voronoi polygron area plot, blue line indicates the obtained data, whereas 99% CI of Monte-Carlo simulation are indicated by red and light blue lines. Representative analysis of two independent experiments is shown. **F** Cluster diameter for Z-138 cells are displayed based on the spatial point distribution analysis. Data (violin plot) are pooled from eight analyzed areas, obtained from two independent experiments per cell line. **G** Cluster stoichiometry analysis of Z-138 cells. Data are pooled mean ± SEM of eight analyzed areas, obtained from two independent experiments per cell line.

Of the tested cell lines, Z-138 stood out as the cell line with the highest level of CXCR4 oligomers (Fig. 2B and S6). To further investigate the link between receptor expression and oligomerization, we tested the effects of genetic manipulations of CXCR4 expression on CXCR4 oligomerization. Doxycycline-inducible expression of CXCR4 enhanced its oligomeric state in RPC1- WM1 (Fig. 2C, S8A) and MEC-1 cells (Fig. S8C), whereas siRNA-mediated silencing of CXCR4 caused a marked reduction of endogenous receptor oligomerization in Z-138 (Fig. 2D, S8B) and Namalwa cells (Fig. S8D). These data effectively demonstrate that CXCR4 expression is an important driver of receptor oligomerization in endogenous systems.

To validate the BRET-based findings of endogenous CXCR4 oligomerization, we employed dSTORM single-molecule imaging (*35*), using AlexaFluor™ 647-conjugated VUN415 (VUN415-AF647), on CHO-K1 (CXCR4^negative^) and Z-138 (CXCR4^high^) cells. To assess the specificity of VUN415- AF647, samples were incubated with an excess of CXCR4 antagonist AMD3100, which is known to displace VUN415 but does not affect CXCR4 oligomerization (*19*) (Fig. S9A, S9B). VUN415 can be competed off by small molecule CXCR4 binding compound AMD3100. Z-138 cells contained specific localized events as demonstrated by their elevated number compared to the corresponding non-specifc localized events in the AMD3100-treated sample (Fig. S9B).

Next, we performed a statistical cluster analysis based on Ripley’s K function, Voronoi segmentation and localization output to analyze the CXCR4 cluster stoichiometry on Z-138 cells. Voronoi segmentation was applied to the obtained spatial distribution patterns of the localized events and compared to random distributions generated by Monte-Carlo simulations (Fig. 2E). Z-138 cells showed significant clustering of CXCR4 receptors (Fig. 2E). The average CXCR4 cluster diameter was 49 ± 39 nm (Fig. 2F). Cluster stoichiometry analysis showed a large population of higher-order CXCR4 oligomers (Fig. 2G). Collectively, these dSTORM findings validate the existence of endogenous CXCR4 oligomers detected by BRET and provide stoichiometric insights into the organization of CXCR4 into multimeric structures in lymphoid cancer cells.

### Pharmacological disruption of endogenous CXCR4 oligomers

In order to investigate a potential function of CXCR4 clusters, we sought to disrupt these clusters and investigate the resulting functional consequences. Previously, we have shown that the minor pocket-binding small molecules IT1t, as well as the N-terminus-binding nanobody VUN401, can disrupt CXCR4 oligomers (*18, 19*). Because two completely different types of molecules (small molecule and nanobody) are able to exert similar effects on CXCR4 oligomers and associated downstream signaling, we wondered whether other CXCR4 ligands displayed a similar mode of action. The identification of other, different, oligomer disruptors would reduce the chance that the phenotypic observations can be contributed to aspecific IT1t and VUN401 effects. First, we evaluated the effects of clinical candidates AMD070 (AMD11070, mavorixafor) and TG-0054 (burixafor) on CXCR4 oligomerization by assessing changes in BRET between Rluc- and YFP-tagged CXCR4 in HEK cells (Fig. 3A) and Spatial-intensity Distribution Analysis (SpiDA, Fig. 3B). In both assays, AMD070, and the controls IT1t and VUN401 reduced the amount of CXCR4 oligomers, whereas TG-0054 did not.

**Figure 3.**
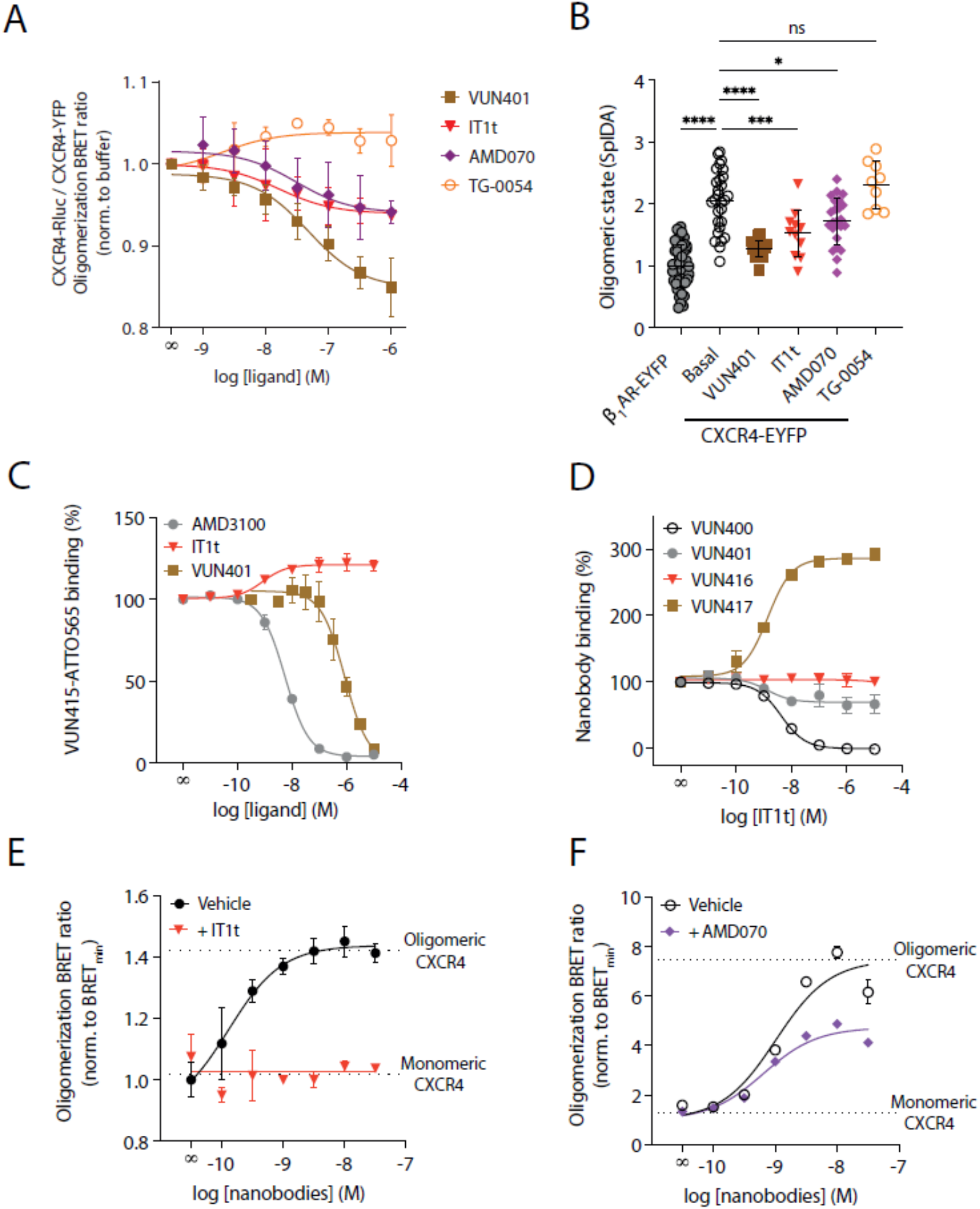
The non-competitive nanobody tool detects IT1t-induced disruption of endogenous CXCR4 oligomers in Z-138 cells. **A** Disruption of CXCR4 oligomerization by indicated concentrations of VUN401, IT1t, AMD070 or TG-0054. BRET values were determined in HEK cells expressing CXCR4-Rluc and CXCR4-YFP and were normalized to vehicle. Data, normalized to the buffer-only condition, are the pooled means from three experiments ± SD. **B** SpIDA analysis of HEK293AD cells expressing monomeric control β1AR-EYFP (gray), vehicle-stimulated CXCR4-EYFP (white) or CXCR4-EYFP after stimulation with VUN401 (10 μM, brown), IT1t (10 μM, red), AMD070 (10 μM, purple) or TG-0054 (10 μM, orange). Data are the mean ± SD, with each data point representing a brightness value from one cell normalized to the monomer control. Data were obtained from three experiments per condition.. **C** Levels of BRET-based measurement of VUN415-ATTO565 (1 nM) displacement by increasing concentrations of indicated CXCR4 antagonists using membrane extracts from NanoLuc-CXCR4-expressing HEK293T cells. **D** Levels of BRET-based measurement of indicated nanobody-ATTO565 (1 nM) displacement by increasing concentrations IT1t using membrane extracts from NanoLuc-CXCR4-expressing HEK293T cells. Data are pooled mean ± SEM of three independent experiments **(C-D)**. **E** VUN416-based BRET measurement of CXCR4 monomerization by 10 μM IT1t in Z-138 cells. **F** VUN415-based BRET measurement of CXCR4 monomerization by 10 μM AMD070 in Z-138 cells. Data are mean ± SD and are representative of three independent experiments, each performed in triplicate.

As IT1t interfered with VUN415 binding to CXCR4 (Fig. 3A, S10), VUN415 can not be used to monitor endogenous modulation of CXCR4 clustering by IT1t. Fortunately, out of a panel of different CXCR4 binding nanobodies , VUN416 binding was unaffected by IT1t (Fig. 3D) and did not modulate CXCR4 oligomerization itself (Table S1). This makes VUN416 a suitable candidate to be engineered into a BRET sensor for the assessment of the effects of IT1t on endogenous CXCR4 oligomers. A mix of VUN416-NanoLuc and VUN416-ATTO565 was able to detect endogenous CXCR4 oligomers in Z- 138 cells, the lymphoid cancer cell line with the highest CXCR4 oligomeric state (Fig. 3E). More importantly, while IT1t did not affect the binding of these probes (Fig. S10B), it completely abolished the oligomer BRET values (Fig. 3E). Fortunately, AMD070 did not affect the CXCR4 binding of oligomer detection nanobody VUN415 (Fig. S10C), allowing the probing of endogenous oligomer disruption by this ligand. Without affecting the binding of the detection nanobodies, AMD070 indeed partially reduced the endogenous CXCR4 oligomers in Z-138 cells (Fig. S3F). This indicates that the oligomer-disrupting activity of IT1t and to a smaller extend AMD070 is also apparent in highly CXCR4 expressin Z-138 cells.

### CXCR4 oligomers drive basal cell migration and anti-apoptotic signaling in MCL cells

To investigate the functional consequences of CXCR4 oligomerization, we first examined the effect of CXCR4 monomerizing small molecule IT1t and nanobody VUN401 on the phosphoproteome of Z-138 cells. Because these cells display the largest CXCR4 oligomerization status, we expect to find the largest changes in protein phosphorylation upon CXCR4-oligomer disruption in these cells. By phospho-enrichment, as described previously (*36*), we identified a total of 15,563 phosphopeptides (purity >80%) (Fig. 4A). This enabled highly sensitive and specific quantifications to be performed across >15,000 phosphosites (localization probability >0.75; Class I), and between cells treated with cluster-disrupting agents and untreated controls (Fig. 4A and S11). Extensive coverage of the phosphoproteome enabled an unbiased and deep characterization of phosphorylation events following CXCR4 cluster disruption, which we subsequently used as a proxy to decipher cellular impact and molecular consequences of interfering with CXCR4 cluster formation.

**Figure 4.**
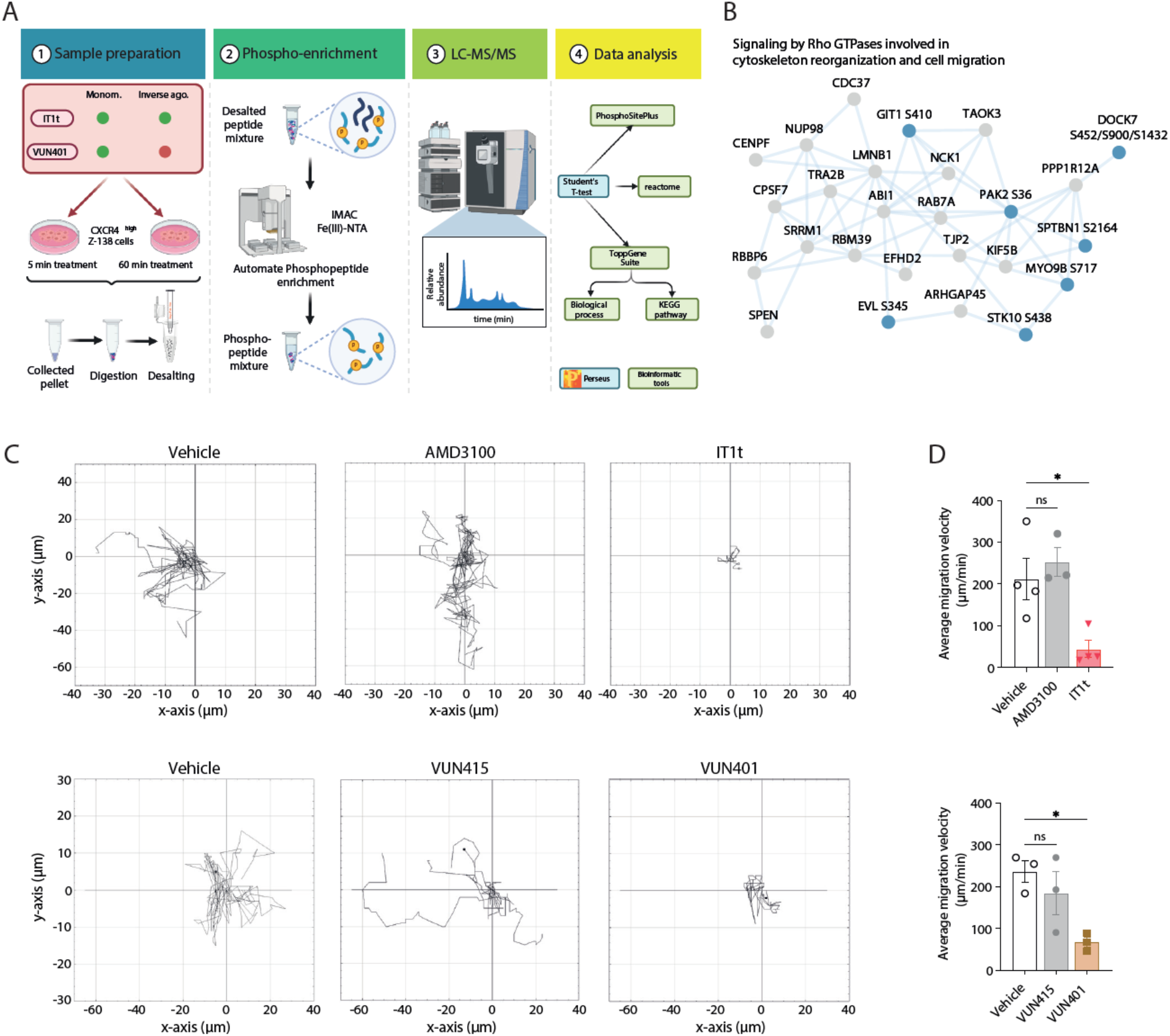
CXCR4 oligomers affect basal cell migration, drive anti-apoptotic signaling and cell viability in MCL cells. **A** Phosphoproteomics study setup and workflow. **B** Protein network of downregulated phosphoproteins in signaling by Rho GTPases involved in cytoskeleton reorganization and cell migration by 60 minutes of 1µM IT1t treatment. The phosphoproteins depicted with bold text are known functional phosphosites. **C** Migration trajectory plots of MCL Z-138 cells for four hours following treatment with 1 μM of AMD3100, IT1t, VUN415 or VUN401. Trajectories are representative of at least three independent experiments. **D** Average velocity following treatment with 1 μM of AMD3100, IT1t, VUN415 or VUN401 derived from average trajectory information. Data are pooled mean ± SEM of at least three independent experiments. * P < 0.05, ** P < 0.01 compared to vehicle, according to unpaired t-tests.

We found that disruption of CXCR4 oligomers changed phosphosites of cell migration mediators, such as the Rac1 effector protein serine/threonine-protein kinase PAK2, GTPase-activating protein DOCK7 and several cytoskeleton rearranging proteins. These phosphoproteins, known to regulate cell migration and reorganizing cytoskeleton, are significantly changed in the signaling pathway regulated by Rho GTPase (Fig. 4B, Fig. S11C). Previously, CXCL12-induced formation of CXCR4 higher-order oligomers has been reported to be essential for sensing chemokine gradients and promoting directed migration of malignant T-cells (*24, 34, 37*). Therefore, we hypothesized that high levels of CXCR4 oligomers might instigate basal signaling towards cell migration. We performed a live-cell imaging experiment, where a consistent proportion of the control-treated cells (5-10%) showed significant basal cell migration during a 4 hour period. Monomerizing ligands IT1t and VUN401 impaired the basal cell migration of this population significantly, whereas non-monomerizing ligands AMD3100 and VUN415 did not (Fig. 4C). When analyzing the trajectories, both IT1t and VUN401 impaired the average migration speed of the highly migratory Z-138 cell population significantly (Fig. 4D). IT1t showed a significant inhibition of the average traveled distance, whereas VUN401 showed a similar trend (Fig. S12). Collectively, these results highlight a role for CXCR4 oligomers in constitutive cell migration, which can be modulated by oligomer disruptors.

To elucidate other CXCR4 clustering-dependent phenotypes, we performed pathway analyses to identify processes similarly affected by IT1t and VUN401 treatment. Of notable interest was the shared positive regulation of the apoptotic process by both IT1t and VUN401 (Fig. 5A), exemplified by the regulation of a large cluster of phosphosites controlling apoptotic events (Fig. 5B). Due to the annotated functions of these phosphorylation sites in regulating apoptosis, and considering the large cluster of coherently regulated phosphosites pointing towards apoptosis, we examined cell viability in more detail. CXCR4 oligomer disruption by IT1t, AMD070 an to a lesser extent VUN401, but not by the non-cluster disrupting molecules AMD3100 and TG-0054, impaired cell viability in the MCL cell lines (Table S3, Fig. S13). Although this data hints towards a protective role for basal CXCR4-oligomer-mediated signaling in these cells, the observed effects were marginal.

**Figure 5.**
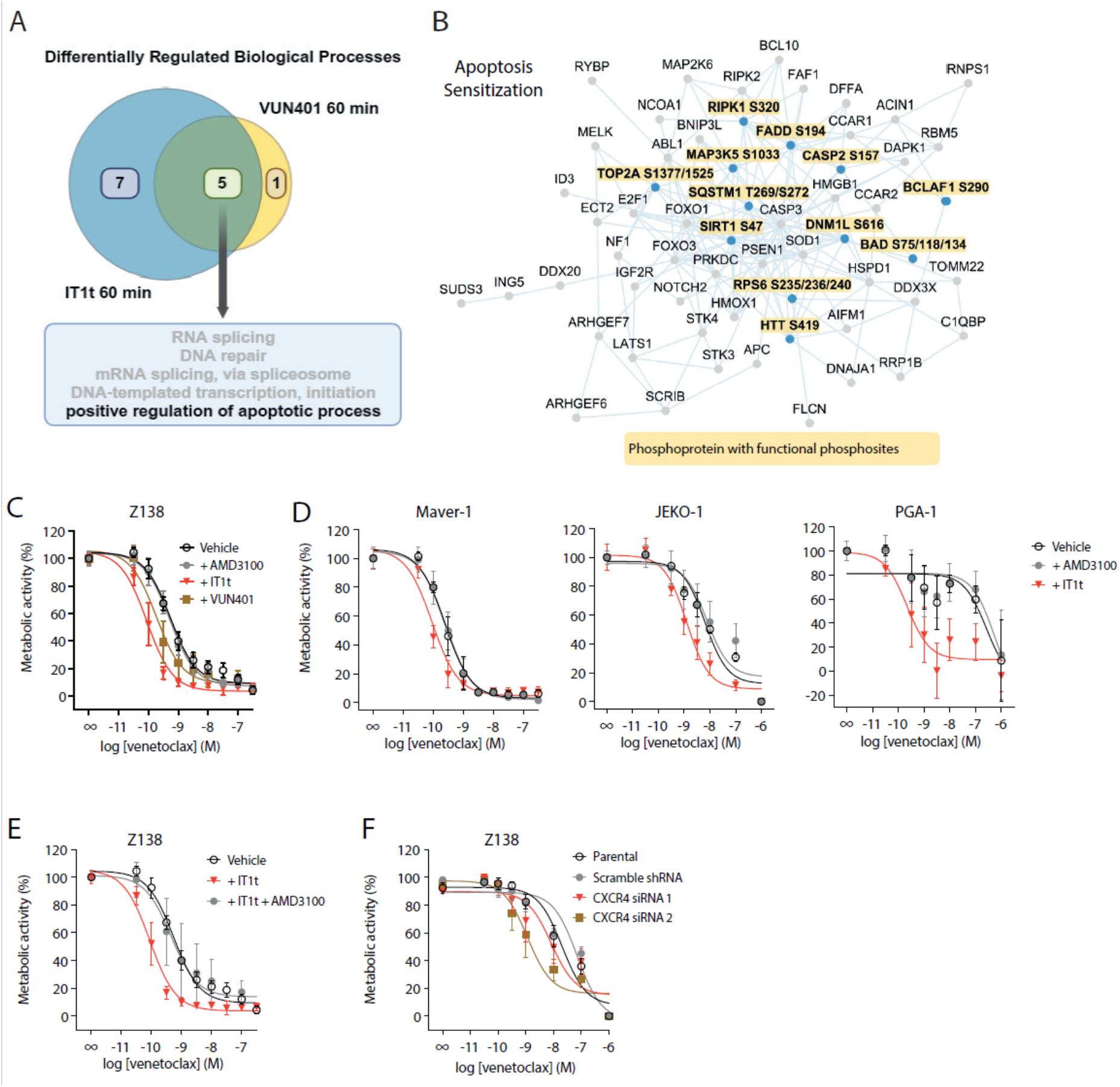
CXCR4-monomerizing ligands sensitize MCL cell line Z-138 to venetoclax-induced apoptosis. **A** Venn diagram depicting overlapping and non-overlapping GO biological processes of 1µM IT1t and VUN401 60 minutes treatment samples. **B** Protein network of downregulated phosphosites in apoptosis sensitization triggered by 60 minutes of 1µM IT1t or VUN401 treatment. The phosphoproteins depicted with bold text, in yellow rectangles are known functional phosphosites. **C** Resazurin-based measurement of metabolic activity in Z-138 MCL cells after 48 h treatment with increasing concentrations venetoclax in absence (vehicle) or presence of 10 μM AMD3100, IT1t or VUN401. Data, normalized to the ‘no venetoclax’ condition, are pooled mean ± SEM of at least three independent experiments, each performed in triplicate. **D** Resazurin-based measurement of metabolic activity in indicated lymphoid cancer cell lines after 48 h treatment with increasing concentrations venetoclax in absence (vehicle) or presence of 10 μM AMD3100 or IT1t. **E** Resazurin-based measurement of metabolic activity in Z-138 cells after 48 h treatment with increasing concentrations of venetoclax in absence (vehicle) or presence of 10 μM IT1t ± 100 μM AMD3100. **F** Resazurin-based measurement of metabolic activity in Z-138 cells upon CXCR4 knockdown and 48 h treatment with increasing concentrations of venetoclax.

Therefore, we tested whether CXCR4-monomerization would increase the sensitivity of Z-138 cells to cell death. venetoclax, a selective Bcl-2 inhibitor, is a cell-death-inducing agent that is approved for CLL and AML patients (*38, 39*). Co-treatment of Z-138 cells with monomerizing ligands IT1t and, to a lesser extent, VUN401 enhanced the sensitivity for venetoclax-induced cell death, as determined by measuring cell metabolic activity with a resazurin assay (Fig. 5C, data normalized to no venetoclax condition for each CXCR4 molecule to emphasize potentiation, Table S2). The effect of IT1t could be blocked by a saturating concentration of the IT1t-competitor and other CXCR4 binder AMD3100 (Fig. 5E), indicating that the effect is CXCR4-specific. Dual concentration-response curves of venetoclax and IT1t revealed a dose-dependent enhancement in the sensitivity of Z-138 cells for venetoclax-induced cell death that saturated at 10 μM IT1t (Fig. S14A). As an alternative approach to disrupt CXCR4 oligomers, we lowered the CXCR4 expresssion by a partial knockdown of CXCR4, which also showed venetoclax sensitization (Fig. S5F). Moreover, the IT1t effect was strongly impaired upon CXCR4 knockdown (Fig. S14B).

In a viability assay by FACS, AMD070 sensitized cells to venetoclax-induced cell death to a lesser smaller extent than IT1t, which corresponds well to its partial oligomer disruption disrupting capability. AMD3100 and the non-monomerizing small molecule TG-0054 did not show sensitization. Sensitization of venetoclax-induced cell death by IT1t was also observed in JEKO-1 cells (Fig. S15B and Table S3). Using the Bliss independence model, a synergistic nature was observed for the enhancement of venetoclax-induced cell death of Z-138 cells by IT1t and, to a lesser extent, VUN401 (Fig. S15A). Pre-incubation with the pan-caspase inhibitor qVD-OPH reduced the observed sensitization, suggesting the involvement of caspases in this process (Fig. S15B). Collectively, these data indicate that CXCR4 clustering promotes anti-apoptotic signaling and associated phenotypes in lymphoid cancer cell lines, which can be targeted using CXCR4-monomerizing ligands.

### Disruption of CXCR4 oligomerization sensitizes to cell death and inhibits spheroid growth in primary CLL and MCL cultures

Finally, we investigated the significance of CXCR4 clusters in patient-derived primary CLL and MCL cells. Using our nanobody-BRET approach, native CXCR4 oligomers could also be detected on primary cells from five CLL patients and two MCL patients. Also on these primary cells, the BRET values that indicate native CXCR4 oligomers, could be almost completely disrupted by IT1t (Fig. 6A), similar to the results obtained with cell lines. More importantly, also in these primary CLL cultures, IT1t, but not the non-monomerizing ligand AMD3100, potentiated the effect of venetoclax (Fig. 6B).

**Figure 6.**
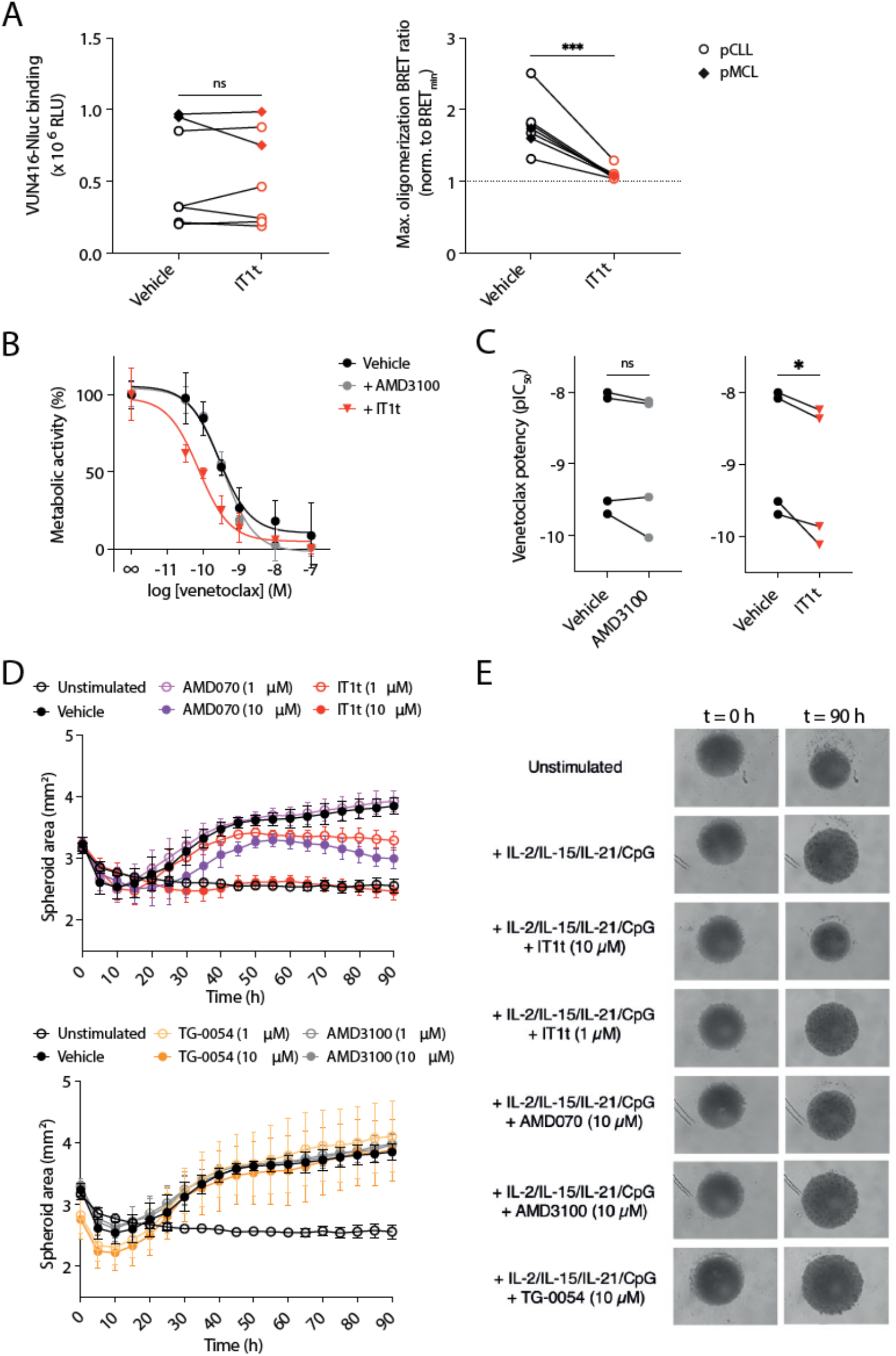
Disruption of CXCR4 oligomerization sensitizes to therapy-induced cell death and inhibits spheroid growth in primary CLL and MCL cultures. **A** Effect of IT1t (10 µM) on VUN416-NanoLuc binding and nanobody-based BRET detection of CXCR4 oligomerization in PBMCs isolated from five CLL and two MCL patients. **B, C** Effects of AMD3100 and IT1t on venetoclax-induced cell death in primary cultures of CLL patients. Full concentration-response curves for one patient **(B)** or ΔpIC50 for four patients **(C)** are shown. **D** Effects of 1 and 10 μM of indicated CXCR4 antagonists on IL-2/IL-15/IL-21/CpG cocktail-induced growth curve in CLL patient-derived spheroid model (41). Data are mean ± SEM of cultures from four (TG-0054) or five individual patients. **E** Effects of indicated CXCR4 antagonists on IL-2/IL-15/IL-21/CpG cocktail-induced growth curve in CLL patient-derived spheroid model after 0 and 90 hours as determined by live-cell imaging (41). Representative images of a culture derived from a single patient are shown. Data are mean ± SEM of cultures from four (TG-0054) or five individual patients. * P < 0.05, ** P < 0.01, **** P < 0.0001, according to unpaired t-tests **(A, C)**.

Given the importance of CXCR4 in lymph node retention (*40*) and the CXCR4 oligomer-driven migration we observed, we tested the effects of CXCR4 ligands in a 3D lymph node-mimicking CLL model derived from patient peripheral blood cells (*41*). CXCR4 monomerizers IT1t and AMD070, and not AMD3100 and TG-0054, inhibited spheroid growth without having cytotoxic effects (Fig. 6D and E, S16, S17A). Compared to AMD070, IT1t inhibited spheroid growth more potently and additionally inhibited the expression of activation marker CD25, of which expression is associated with poor desease outcome (Fig. S17B). Taken together, our data indicate that CXCR4 oligomers also contribute to pro-survival signaling in CLL patient-derived cultures and that specific disruption of such oligomers is a promising therapeutic outlook.

## Discussion

Despite major improvements in the treatment of B cell lymphoid neoplasms, many patients still experience relapsing disease that becomes more aggressive with each recurrence, characterized by acquired resistance and impaired clinical outcome. Hence, there is a need for novel therapeutic interventions, preferably targeting other signaling pathways with other modes of action. In this work, we have uncovered that CXCR4 oligomers exist on many B cell lymphoid cancer cells. Also, our data indicates that such oligomers can induce oncogenic signaling and that inhibiting this signaling through oligomer disruption represents a novel therapeutic strategy for the treatment of CLL, MCL and other B cell lymphoid neoplasms.

High expression of CXCR4 is known to correlate with tumor growth, invasion, relapse and therapeutic resistance (*3*). For example, in CLL patients, high CXCR4 expression is associated with reduced progression-free survival (*42*). The signaling output of GPCRs, such as CXCR4, is generally evaluated in the context of agonist stimulation. For instance, there is ample evidence that supports the cancerous role of CXCL12-mediated migration and pro-survival signaling (*3, 4, 6*). However, apart from agonist-driven receptor activation, constitutive GPCR signaling exists and, in the case of CXCR4 signaling, has been reported to promote the growth and survival of acute myeloid leukemia *in vivo* (*43*). Moreover, colon cancer cells require CXCR4 expression but not ligand-induced signaling capacity for chemotherapy resistance (*44*). Our findings indicate that the high oligomeric state of CXCR4 in lymphoid cancer cells induces constitutive pro-survival signaling and basal migration. Similarly, the constitutive activity of breast tumor kinase and the adhesion GPCR GPR64 accelerates cell migration and thereby contributes to tumorigenesis (*45, 46*). Constitutive receptor oligomerization-driven signaling has been observed for other GPCRs, as exemplified by the requirement of CCR7 oligomer formation for the interaction with and activation of tyrosine kinase Src (*47*). The CXCR4 oligomerization-driven basal cell migration, likely contributing to the invasive and metastatic properties associated with this receptor, might be particularly important for cells that are not exposed to a CXCL12 gradient.

Most studies focusing on GPCR oligomerization and modulation were performed in heterologous expression systems, which may differ from endogenous systems. For instance, the M_1_R antagonist pirenzepine promotes oligomerization of transfected M_1_R receptors but prevents the formation of endogenous oligomers (*48–50*). Employing our nanobody-based tools, we report the pharmacological disruption of endogenous CXCR4 oligomers for the first time. AMD070 partially disrupted CXCR4 oligomers in lymphoid cancer cells, while IT1t appeared to induce a fully monomeric receptor state. This was slightly different in heterologous expression systems, where equal oligomer-disrupting efficacies were observed for these two small molecules. These data highlight the importance of studying oligomerization in a native context.

We have tested several small molecules for their ability to disrupt CXCR4 oligomers. Of these, AMD3100, an antagonist lacking oligomer-disrupting properties, is used clinically for hematopoietic stem cell mobilization (*51*). In addition, the partial cluster disruptor AMD070 has recently been approved for the treatment of WHIM syndrome, demonstrating that antagonizing CXCR4 is clinically safe (*52*). In contrast to AMD3100 and TG-0054, we found that AMD070, IT1t, and nanobody VUN401 disrupted CXCR4 oligomers and inhibited downstream signaling towards cell survival and migration. The effects of IT1t could theoretically also be attributed to its ability to inhibit basal CXCR4-mediated Gα_i/o_ signaling. However, VUN401 and AMD070 do not inhibit basal CXCR4-mediated Gα_i/o_ signaling (*19*) and IT1t and VUN401 showed similar effects on anti-apoptotic and cell migration signaling networks in our studies. This suggests inhibition of CXCR4 oligomerization to be the underlying mechanism rather than inhibition of basal Gα_i/o_ activation.

In our venetoclax sensitization experiments and CLL patient-derived 3D spheroid model, the full monomerizing ligand IT1t was more potent and efficacious than the partial cluster disruptor AMD070. This implies that the extent to which CXCR4 oligomers can be disrupted may impact the therapeutic outcome, at least in lymphoid neoplasms. In these pathologies, high efficacy cluster disruptors would be the most attractive candidates for the therapeutic targeting of CXCR4. As expression is a primary driver of CXCR4 oligomerization, pharmacological intervention with cluster disruptors would specifically target malignant cells that overexpress CXCR4. This would create an added layer of selectivity for targeted therapy in cancer. Our approach could be expanded to other GPCRs, like P2Y2 receptors, which are highly expressed in pancreatic cancer and form clusters that can be pharmacologcically disrupted (*53*).

The pathological role of CXCR4 oligomerization described in this study may extend beyond the context of lymphoid neoplasms. Previously, IT1t but not AMD3100 was shown to inhibit TLR7- mediated type I interferon signaling in plasmacytoid dendritic cells from systemic lupus erythematosus patients (*54*). Moreover, IT1t inhibited early metastases in an *in vivo* breast cancer zebrafish model (*55*). Although future studies are required to investigate whether these phenotypes can be (fully) ascribed to the oligomer-disruptive effects of IT1t, this added capacity may prove beneficial over inhibiting CXCL12-induced Gα_i/o_ signaling by antagonizing compounds like AMD3100. The therapeutic potential of inhibiting CXCR4 oligomer-mediated basal signaling might also be extended towards other CXCR4 overexpressing cancer types. Since disruption of CXCR4-oligomers inhibited multiple hallmarks, it is not unlikely that cluster disruption can result in the potentiation of other commonly used cell-death-inducing agents.

Taken together, this study demonstrates the existence of native CXCR4 oligomers in lymphoid neoplasms and CXCR4 oligomer-driven signaling with pathophysiological importance. Selective targeting of CXCR4 clustering in lymphoid neoplasms and other cancers may have therapeutic potential on its own or by potentiating other therapeutics.

## Methods

### DNA constructs and molecular cloning

The pcDEF3 vector was a gift from Langer (*56*). cDNA encoding the BRET-based cAMP biosensor His-CAMYEL pcDNA3.1(L) was purchased from ATCC (#ATCC-MBA-277). pLKO.1 puro CXCR4 siRNA-1 and siRNA-2 were gifts from Bob Weinberg (Addgene plasmids #12271 and #12272). plKO.1 scramble shRNA was a gift from David Sabatini (Addgene plasmid #1864). Myc-CXCR4-Rluc pIRES, HA-CXCR4-YFP pIRES (*17*), HA-CXCR4 WT pcDEF3 (*57*), HA-CXCR4-N119S pcDEF3 (Bergkamp and Perez Almeria et al*, manuscript in preparation*), NanoLuc-CXCR4 pcDNA3.1 (*58*) and CXCR4- EYFP pcDNA3 (*19*) were described previously. HA-CXCR4 pLenti6.3/TO/V5-DEST was generated by exchanging US28 for CXCR4 in the previously described HA-US28 pLenti6.3/TO/V5-DEST plasmid (*59, 60*). HA-CXCR4 pEUI was generated by exchanging VUN103-FLAG for HA-CXCR4 in the previously described VUN103-FLAG pEUI plasmid (*61*).

### Patient material

After written informed consent, patient blood samples were obtained during diagnostic or follow-up procedures at the Departments of Hematology and Pathology of the Academic Medical Center Amsterdam. This study was approved by the AMC Ethical Review Biobank Board under the number METC 2013/159 and conducted in accordance with the Declaration of Helsinki. Peripheral blood mononuclear cells (PBMCs) of patients with CLL, obtained after Ficoll density gradient centrifugation (Pharmacia Biotech), were cryopreserved and stored as previously described (*62*). On the day of the experiment, the PBMCs were thawed in a water bath at 37°C. Thawing medium, consisting of Iscove’s Modified Dulbecco’s Medium (IMDM, Gibco) supplemented with 100 units of penicillin, 100 g/mL streptomycin (P/S, Gibco) and 20% (v/v) Fetal Bovine Serum (FBS, Bodinco), was added and cells were rested in the dark for 20 min at RT. Next, thawing medium was removed and cells were washed by centrifuging for 5 min at 300 x g with the deacceleration rate set at 7. Cells were then resuspended in assay buffer, consisting of IMDM supplemented with 10% FBS and 1% P/S. Cells were counted and a viability of ≥ 70% was ensured by conducting a trypan blue staining using a LUNA-II™ automated brightfield cell counter (Logos Biosystems).

### Cell lines and cell culture

Human embryonic kidney 293T (HEK293T) and CHO-K1 cells were obtained from American Type Culture Collection (ATCC). MEC-1, PGA-1, L363, CCRF-CEM, Jeko-1, CII, Namalwa, Maver-1 and Z-138 were described previously (*63*). RPCI-WM1 and TMD8 were kindly provided to Marcel Spaargaren by Dr. S.P. Treon and Dr. G. Lenz, respectively. HEK293T cells were cultured in Dulbecco’s Modified Eagle’s Medium (DMEM, Gibco) supplemented with 10% FBS and 1% P/S. CHO-K1 cells were cultured in DMEM/F-12, supplemented with 10% FBS and 1% P/S. MEC-1, RPCI-WM1, TMD- 8, PGA-1, L363, CCRF-CEM, Jeko-1, CII, Namalwa, Maver-1 and Z-138 cells were cultured in IMDM supplemented with 10% FBS and 1% P/S. One day prior to experiments, suspension cells were prepared at a concentration of 1 x 10^6^ cells/mL and adherent cells were maintained in culture under non-confluent conditions. On the day of the experiment, cells were recounted and viability of ≥ 90% was ensured by conducting a trypan blue staining using a LUNA-II™ automated brightfield cell counter.

### Nanobody generation and production

Previously described nanobodies were cloned into the pMEK222 bacterial expression vector with C- terminal FLAG-6xHis tag (*27, 64, 65*). VUN415-NanoLuciferase (VUN415-NanoLuc) and VUN416- NanoLuc were generated by subcloning VUN415 and VUN416 into a modified version of the pMEK222 vector, with a C-terminal upper Hinge linker–NanoLuciferase–6xHis tag (*66*). BL21 Codon+ bacteria transformed with these pMEK222 plasmids were grown O/N in 10 mL of 2xYT medium, supplemented with glucose (2%) and ampicillin (1 μg/mL). Next day, this O/N culture was inoculated (1:100) in Terrific Broth with ampicillin (1 μg/mL). After bacteria grew at 37 °C to OD600 of 0.5, nanobody production was initiated by adding isopropyl β-D-1-thiogalactopyranoside (1 mM) and incubation took place for 4 h at 37 °C. After centrifugation, pellets were frozen O/N at -20 °C. After thawing and dissolving the pellet in phosphate-buffered saline (PBS, pH 7.4), periplasmic extracts were incubated head-over-head for 1.5 h at 4 °C. Nanobodies were purified from the periplasm using immobilized affinity chromatography (IMAC) via 6x-His tags. Nanobodies bound to ROTI®-Garose cobalt agarose beads (Carl Roth) were eluted with 150 mM imidazole (Sigma-Aldrich). Afterwards, the buffer of nanobody eluates was exchanged for PBS by O/N dialysis using Snakeskin Dialysis Tubing (Thermo Fisher Scientific). Dialyzed fractions were combined and stored at -20 °C until experiments.

### Fluorescent labeling of nanobodies

The labeling of CXCR4 nanobodies with ATTO565 fluorescent dyes (ATTO-TEC, #AD565-41, #AD565-31) using thiol-maleimide coupling and N-hydroxy-succinimide (NHS) chemistry was described previously (*58*). Nanobodies containing an unpaired cysteine in the C-terminal tag used for fluorophore thiol-maleimide conjugation were provided by QVQ (Utrecht, the Netherlands). UV-VIS spectrometry was performed to ensure degree of labeling (DOL) > 0.5. Free dye of <5% was assessed by SDS-PAGE, followed by a fluorescence scan using an Odyssey imager (LI-COR, at suboptimal wavelength to prevent detector saturation) or Azure400 imager (Azure Biosystems, at 524 nm excitation). In a similar fashion, VUN415-Cys was conjugated using thiol-maleimide coupling with an excess of Alexa Fluor 647 C2-maleimide (Invitrogen, A20347), to ensure a DOL of 1.

### Transfection HEK293T cells

HEK293T cells were transfected with a total of 1 μg DNA and 6 μg 25 kDa linear polyethyleimine (PEI, Polysciences Inc.) in 150 mM NaCl solution per 1 × 10^6^ cells. DNA encoding receptors and biosensors was, if necessary, supplemented with empty pcDEF3 to obtain a total DNA amount of 1 μg. The DNA-PEI mixture was vortexed for 3 s and incubated for 15 min at room temperature (RT). HEK293T cells were detached with Trypsin (Gibco) and resuspended in DMEM. The HEK293T cell suspension was added to the DNA-PEI mixture and cells were seeded at 3.5 × 10^4^ per well in white flat-bottom 96-well plates (Greiner Bio-One).

### Receptor oligomerization CXCR4-Rluc and CXCR4-YFP

For receptor oligomerization experiments using tagged receptors, HEK293T cells were transfected with 40 ng Myc-CXCR4-Rluc and 400 ng HA-CXCR4-YFP. After 48 h incubation, cells were washed once using PBS and maintained in Hank’s Buffered Saline Solution (HBSS), supplemented with 0.1% BSA, 1 mM MgCl2 and 2 mM CaCl2. Cells were stimulated with increasing concentrations of CXCL12, small molecules or nanobodies for 15 min before BRET measurements. After incubating cells for 10 min with 5 μM coelenterazine-h substrate (Promega), bioluminescence was measured at 535/30 nm and 475/30 nm using a PHERAstar plate reader (BMG). BRET signals were determined as the ratio of luminescence in the acceptor channel divided by the donor channel. The ligand-promoted BRET signal was calculated by dividing the pre-read-normalized BRET values of each ligand concentration by the BRET ratio obtained for the vehicle condition.

### CAMYEL constitutively active CXCR4

To assess potential inverse agonism of ligands on basal Gα_i/o_ activation, HEK293T cells were transfected with 500 ng constitutively active CXCR4 mutant (HA-CXCR4 N119S) and 500 ng CAMYEL. After 48 h incubation, cells were washed once using PBS and maintained in HBSS, supplemented with 0.1% BSA, 1 mM MgCl2 and 2 mM CaCl2. After 20 min stimulation with 100 nM CXCL12, 10 μM small molecules, 1 μM VUN415 and 15 min stimulation with 1 μM forskolin (i.e. adenylyl cyclase activator), BRET measurements were performed. After incubating cells for 10 min with 5 μM coelenterazine-h substrate, bioluminescence was measured at 535/30 nm and 475/30 nm using a PHERAstar plate reader.

### Oligomer detection using nanobody-based BRET in transfected HEK293T cells

To detect nanobody-based oligomerization BRET, HEK293T cells were transfected with 500 ng HA- CXCR4 pcDEF3 or HA-CXCR4 pEUI. For FKBP experiments, HEK293T cells were transfected with 2 ng HA-CXCR4 or 2 ng HA-CXCR4-FKBP. For HA-CXCR4 pEUI, increasing concentrations tebufenozide (Sigma-Aldrich) were added into the culture medium 6 h post-transfection. After 48 h, cells were washed once with PBS. In the case of FKBP experiments, cells were treated with or without 1 μM AP20187 for 1 h. Subsequently, increasing equimolar concentrations of VUN415-NanoLuc and VUN415-ATTO565 or a constant concentration of detection nanobodies (31.6 nM) with different donor to acceptor ratios in assay buffer (HBSS, supplemented with 0.1% BSA, 1 mM MgCl2 and 2 mM CaCl2) were added to the cells. After incubation for 2 h at RT, cells were washed twice with PBS and assay buffer was added. Subsequently, fluorescence of fluorescently labeled nanobodies was measured using a CLARIOstar plate reader at 563/30 nm excitation and 592/30 nm emission. After addition of 15 μM furimazine substrate (NanoGlo, Promega), luminescence was measured using a PHERAstar plate reader with 610 nm/LP and 460/80 nm filters until the luminescence signal stabilized.

### ELISA for surface expression of ecdysone-inducible CXCR4

In parallel with the BRET experiment described before, 3.5 × 10^4^ transfected HEK293T cells were seeded in a transparent flat-bottom 96-well plate (Greiner Bio-One). Increasing concentrations of tebufenozide were added into the culture medium 6 h post-transfection. After 48 h, cells were fixated using 4% paraformaldehyde (PFA) in PBS and plates washed with PBS. Subsequently, blocking was performed with 2% (w/v) skimmed milk in PBS for 1 h at RT. Antibody incubations were also performed using this blocking buffer. CXCR4 expression was detected with the monoclonal mouse anti-CXCR4 antibody 12G5 (1:1000, Thermo Fisher Scientific, #35-8800) and horseradish peroxidase (HRP)- conjugated goat-anti-mouse antibody (1:2000, Bio-Rad, #1706516). Incubations with these antibodies were performed for 1 h at RT. Wells were washed three times with PBS between all incubation steps. Binding was determined with 1-step Ultra TMB-ELISA substrate (Thermo Fisher Scientific), and the reaction was stopped with 1M H2SO4. Optical density was measured at 450 nm using a CLARIOstar plate reader.

### Membrane extract preparation

Two million HEK293T cells were plated in a 10 cm^2^ dish (Greiner Bio-One). The next day, cells were transfected with 250 ng NanoLuc-CXCR4, supplemented to a total of 5 μg DNA with empty pcDEF3 vector, and 30 μg PEI in 150 mM NaCl solution. The DNA-PEI mixture was vortexed for 3 s and incubated for 15 min at RT. Subsequently, the mixture was added dropwise to the adherent HEK293T cells. Protein expression was allowed to proceed for 48 h. Media was then removed and cells were washed once with cold PBS. Next, cells were detached and resuspended in cold PBS. Cells were centrifuged at 1500 × *g* at 4 °C, resuspended in cold PBS, and again centrifuged at 1500 × *g* at 4 °C. The pellet was resuspended in membrane buffer (15 mM Tris-Cl, 0.3 mM EDTA, 2 mM MgCl2, pH 7.5) and disrupted by the homogenizer Potter-Elvehjem at 1200 rpm. Next, membranes were freeze-thawed using liquid nitrogen, pelleted by ultracentrifugation (25 min, 40000 × *g*, 4°C), carefully washed with Tris-Sucrose buffer (20 mM Tris, 250 mM Sucrose, pH = 7.4 at 4°C) and resuspended in Tris-Sucrose buffer. The membranes were homogenized using a 23G needle (10 strokes), aliquoted, snap-frozen using liquid nitrogen and protein concentrations were determined using a bicinchoninic acid assay (Pierce™ BCA Protein Assay; Thermo Fisher Scientific). Subsquently, the membranes were stored at −80°C until use in NanoBRET assays.

### Displacement of fluorescent nanobodies and CXCL12

Approximately 0.25 μg per well of membrane extracts from NanoLuc-CXCR4-expressing HEK293T cells was added to a white flat-bottom 96-well plate. Subsequently, increasing concentrations of unlabeled ligands in HBSS, supplemented with 0.1% BSA, 1 mM MgCl2 and 2 mM CaCl2. The plate was spun down and incubated for 30 min at RT. Next, 316 pM nanobody-ATTO565 or 10 nM CXCL12- AZ647 (Protein Foundry) was added and incubated for 1 h at RT. Next, 15 μM furimazine substrate was added and luminescence was measured using a PHERAstar plate reader with 610 nm/LP and 460/80 nm filters until the luminescence signal stabilized.

### Flow cytometry for CXCR4 surface expression determination

For each sample, 5 x 10^5^ cells were washed with ice-cold FACS buffer (0.5% BSA (PanReac AppliChem, A6588,0100) in PBS) and resuspended in ice-cold FACS buffer containing 3 µg/mL mouse anti-CXCR4 antibody 12G5 (Thermo Fisher Scientific, 35-8800) in polypropylene 5-mL tubes (Falcon). Following incubation on ice for 1 h, samples were washed three times with excess ice-cold FACS buffer to remove unbound antibody. Subsequently, samples were resuspended in ice-cold FACS buffer containing 2 µg/mL goat anti-mouse IgG (H+L) AlexaFluor™ 488 (Thermo Fisher Scientific, A-11001). After incubation and washing as described before, samples were resuspended in ice-cold FACS buffer. Subsequently, samples were analyzed utilizing an Attune Nxt Flow Cytometer (Thermo Fisher Scientific) at the AUMC Microscopy Cytometry Core Facility (MCCF), with flow rates not exceeding 500 µL/min. Sample analysis was conducted using FlowJo version 10 (BD Biosciences) to determine CXCR4 surface expression levels.

### Oligomer detection using nanobody-based BRET in lymphoid cancer cell lines

1 × 10^6^ lymphiod cancer cells were seeded in a white flat-bottom 96-well plate. In case of small molecule disruption, 10 μM of IT1t or AMD070 was added to the cells. Subsequently, cells were stimulated with increasing equimolar concentrations of VUN415/VUN416-NanoLuc and VUN415/VUN416-ATTO565 or ITGB1-Nb-HL555 (QvQ) in assay buffer (HBSS, supplemented with 0.1% BSA, 1 mM MgCl2 and 2 mM CaCl2). For oligomer detection on PBMCs derived from CLL patients, 31.6 nM of VUN416- NanoLuc/-ATTO565 detection nanobodies were added with a ATTO565: NanoLuc ratio of 0.25 (BRET_min_) or 19 (BRET_max_). After incubation for 2 h at RT, cells were washed twice with PBS and assay buffer was added. Subsequently, fluorescence of fluorescently labeled nanobodies was measured using a CLARIOstar plate reader at 563/30 nm excitation and 592/30 nm emission. After addition of 15 μM furimazine substrate, luminescence was measured using a PHERAstar plate reader with 610 nm/LP and 460/80 nm filters until the luminescence signal stabilized.

### Lentivirus production and transduction

MEC-1 and RPCI-WM1 cell lines with inducible HA-CXCR4 expression and Namalwa and Z-138 cell lines with constitutive siRNA CXCR4 or scramble shRNA expression were generated by lentiviral transduction, as previously described (*59, 67*). Briefly, lentivirus was produced for 48 h after co-transfecting four dishes of 2 x 10^6^ HEK293T cells with HA-CXCR4 pLenti6.3/To/V5-DEST, pLKO.1 puro CXCR4 siRNA-1/2 or plKO.1 scramble shRNA together with pRSV-REV, pMDLg/pRRE and pMD2.g packaging vectors, using PEI as transfection reagent. Lentivirus solution from four dishes was pooled, cleared by centrifugation for 10 min at 500 x g and filter-sterilized. Subsequently, lentivirus was ultracentrifuged for 1 h at 70000 x g and supernatant was discarded until approximately 1 mL concentrated lentivirus solution was remaining. This lentivirus solution was then aliquoted and stored at -80C until lentiviral transduction. At the day of lentiviral transduction, 100 µL of concentrated lentivirus solution was added to 1 x 10^6^ cells in 1 mL. Subsequently, cells were incubated for three days before addition of the appropriate antibacterial selection agent. Knockdown efficiency and enhanced CXCR4 surface expression in the different cell lines was validated by determining CXCR4 surface expression levels as described before. CXCR4 expression in the doxycycline-inducible cell lines was induced using 1 µg/mL doxycycline (Sigma-Aldrich).

### dSTORM microscopy

#### Sample preparation

RPCI-WM1, Z-138 and CHO-K1 cells were fixated using 4% paraformaldehyde (PFA) in PBS for 15 min at 37°C. Next, cells were washed once and resuspended in FACS buffer (0.05% BSA in PBS). The fixated cells were then subjected to staining with VUN415-AF647 at RT for 1 h. Unbound VUN415- AF647 was removed through a series of three consecutive washing steps using FACS buffer. 1 x 10^6^ cells were added to a poly-l-lysine (Sigma)-coated coverslip (VWR) in a 6-well plate (Greiner Bio-One). The coverslip was subjected to centrifugation in the 6-well plate at 500 x g for 15 min using a plate centrifuge (Eppendorf). Following this, the samples were stored in suspension in a dark environment at 4°C until the time of readout.

Before imaging, samples were mounted in oxygen scavenger-containing Glox-buffer to facilitate blinking conditions. Glox-buffer was prepared as described previously (*68*). Briefly, the following stock solutions were prepared and stored at -80°C: 1M Cysteamine (MEA) in 250 mM (Sigma, in 250 mM HCl), 70 mg/mL glucose-oxidase (Sigma-Aldrich) and 4 mg/mL catalase (Sigma-Aldrich). When mounting the sample, the final buffer was prepared freshly by diluting stock solutions MEA, glucose-oxidase plus catalase and glucose solution in 50 mM Tris pH 8.0 (final concentrations: 100 mM MEA, 700 μg/mL glucose oxidase, 40 μg/mL catalase, 5% w/v glucose). To prevent oxygen from entering the sample during imaging, coverslips were mounted on cavity slides (Sigma-Aldrich) filled with imaging buffer. By removing surplus buffer from the sides of the coverslip, a vacuum seal was created.

#### Imaging

Imaging was performed on a Ti-E microscope (Nikon) equipped with a 100x Apo TIRF oil immersion objective (NA. 1.49) and Perfect Focus System 3 (Nikon). A Lighthub-6 laser combiner (Omicron) containing a 647 nm laser (LuxX 140 mW, Omicron) and a 405 nm diode laser (Power technology, 15 mW) together with optics allowing for a tunable angle of incidence were used for excitation. Illumination was adjusted for (pseudo-) total internal reflection fluorescence (TIRF) microscopy to remove out-of-focus signal. To separate emission light from excitation light, a quad-band polychroic mirror (ZT405/488/561/640rpc, Chroma) and a quad-band emission filter (ZET405/488/561/640m, Chroma) were used. Detection of the emission signal was done using a Hamamatsu Flash 4.0v2 sCMOS camera. Image stacks were acquired with a 30 ms exposure time, 50-100% laser power of 647 laser, 3-5% laser power of the 405 laser which was increased during imaging, and 5000 images per field of view. Components were controlled using MicroManager (*69*).

#### Data analysis

Acquired stacks were analyzed using v.1.2.1 of a custom ImageJ plugin called DoM (Detection of Molecules) (https://github.com/ekatrukha/DoM_Utrecht), as previously described (*68*). Briefly, each image in an acquired stack was convoluted with a two-dimensional Mexican hat kernel which matches the microscope’s point spread function (PSF) size. The resulting intensity histogram was utilized to create a thresholded mask that was used to calculate the centroids on the original image. These centroids were used as initial values to perform unweighted nonlinear least squares fitting with a Levenberg-Marquardt algorithm to an asymmetric two-dimensional Gaussian PSF, allowing for the sub-pixel localization of particles. The acquired localization output by DoM was imported into the application ClusterViSu (https://github.com/andronovl/SharpViSu) that conducts a statistical cluster-analysis based on Ripley’s K-function and Voronoi segmentation, as previously described (*70*). Eight areas per sample were examined for the RPCI-WM1 and Z-138 samples with an average area of 35 ± 10 µm^2^ and 27 ± 7 µm^2^ respectively. Four areas per sample were examined for the CHO-K1 and non-specificity control samples (i.e. displacement with an excess of CXCR4 antagonist AMD3100). Selected areas did not overlap or came in contact with the edges of the corresponding analyzed cell. Ripley’s K-function was calculated and Voronoi segmentation conducted for the indicated areas and localization distributions compared to a random distribution based on a similar surface area and number of localized points by conducting Monte-Carlo simulations. Segmentation was conducted subsequently by automatic thresholding of the cluster map. Quantitative output, including cluster area, diameter and stoichiometry, were determined.

### Spatial-intensity Distribution Analysis (SpiDA)

For SpiDA analysis, 2.5 × 10^5^ HEK293AD cells were grown on glass coverslips in six-well plates. Next day, cells were transfected with 600 ng of CXCR4-EYFP using Effectene transfection reagent (Qiagen) according to the manufacturer’s protocol. The next, the coverslip was loaded into the Attofluor imaging chamber (Thermo Fisher Scientific). Prior to imaging, cells were stimulated for 30 min with 10 μM IT1t, AMD0070 or TG-0054 in HBSS supplemented with 0.1% BSA. Imaging was performed using a commercial laser-scanning confocal microscope (Leica SP8) equipped with a 63×/1.40 NA oil immersion objective, a white light laser (WLL), and photon counting hybrid detectors. For excitation, 514 nm lines of the WLL were used, and for the detection of EYFP, emission bands of 520 nm to 600 nm were used. Images were acquired using 15% laser power. The image format was xy and image size was set to 512 × 512 pixels with 50-nm pixel size. For image analysis, the open-source custom-made code (https://github.com/PaoloAnnibale/MolecularBrightness) was loaded onto the Igor Pro software (WaveMetrics). Polygonal region of interest (ROI) selection was performed to avoid regions with non-homogenous fluorescence distribution (e.g. membrane raffles, clusters).

### Phosphoproteomics

#### Sample preparation

Cell pellets were lysed in 8 M urea with 50 mM ammonium bicarbonate (pH 8, Sigma-Aldrich) with 1× Protease inhibitor cocktail EDTA (Roche) and 1× PhosSTOP (Roche). Sonication was performed with a Bioruptor (Diagenode) sonicator for 5 cycles (30 s on, 30 s off) at 4 °C. The lysate was spun down for 1 h at 14,000 rpm at 16 °C to pellet cell debris and DNA. Protein concentration was determined by a microplate Bradford assay (Sigma-Aldrich). 1 mg aliquot of each sample was taken for further digestion and phosphopeptide enrichment.

Protein samples were reduced in 10 mM dithiothreitol (DTT, Sigma-Aldrich) at 20 °C for 60 min, and alkylated in the dark with 20 mM iodoacetamide (IAA, Sigma-Aldrich) at 20 °C for 30 min.

An additional final concentration of 10 mM DTT was added to quench the excess IAA. 50 mM ammonium bicarbonate was used to dilute to reach a final concentration of 2 M Urea. The alkylated proteins were sequentially digested using Lys-C (Wako) and trypsin (Sigma-Aldrich) at a 1:75 enzyme-to-protein ratio, and carried out at 37 °C. The Lys-C digestion lasted for 4 h. After which, 50 mM ammonium bicarbonate was used to dilute the samples to a final concentration of 2 M urea, and followed by overnight trypsin digestion with trypsin was performed overnight. 3% formic acid was used to quench the digestion, and digested peptides were desalted by Sep-Pak C18 1 cc Vac cartridges (Waters), dried using a vacuum centrifuge, and stored at -80 °C for further use.

#### Automated Fe^3+^-IMAC phosphopeptide enrichment

Phosphopeptides were enriched by using Fe(III)-NTA 5 μlL (Agilent Technologies) in an automated AssayMAP Bravo Platform (Agilent Technologies). Fe(III)-NTA (nitrilotriacetic acid) cartridges were first primed with 250 μL of priming buffer (99% acetonitrile (ACN), 0.1% TFA) at a flow rate of 100 μL/min and equilibrated with 250 μL of loading buffer (80% ACN, 0.1% TFA) at a flow rate of 50 μL/min. Dried peptides were dissolved in 210 μL of loading buffer and centrifuged at 14000 rpm for 10 min. Samples were then loaded at a flow rate of 3 μL/min onto the cartridge, the flowthrough was collected into a separate plate. Cartridges were washed with 250 μL of loading buffer at a flow rate of 20 μL/min, and the phosphopeptides were eluted with 50 μL of 10% ammonia at a flow rate of 5 μL/min directly into 50 μL of 10% formic acid. Flowthroughs and elutions were dried and injected directly on a liquid chromatography-coupled mass spectrometer.

#### LC–MS/MS analyses

The phosphoproteome measurement was performed on an Orbitrap Exploris 480 mass spectrometer (Thermo Fisher Scientific) coupled with an UltiMate 3000 UHPLC system (Thermo Fisher Scientific) fitted with a µ-precolumn (C18 PepMap100, 5 µm, 100 Å, 5 mm × 300 µm; Thermo Fisher Scientific). Samples were analyzed in triplicates and separated on an analytical column (Poroshell 120 EC-C18, 2.7 µm, 50 cm × 75 µm, Agilent Technologies) with a 115-min gradient. Peptides were first eluted at a constant flow rate of 300 nl/min using 9 to 36% solvent B (0.1% v/v formic acid in 80% acetonitrile) over 97 min, raised to 99% in 3 min, then held for 3 min and equilibrated in 9% B for 1 min. The mass spectrometer was operated in data-dependent mode. Electrospray ionization was performed at a 2.1 kV static spray voltage; the temperature of the ion transfer tube was set to 275 °C, and the RF lens voltage was set to 55%. Full scan MS spectra from the m/z range of 375-1600 were acquired at a resolution of 60,000 after accumulating to the ‘Standard’ pre-set automated gain control (AGC) target. Higher energy collision dissociation (HCD) was performed with 35% normalized collision energy (NCE), at an orbitrap resolution of 30,000. Dynamic exclusion time was set to 90 s and a 0.7 m/z isolation window was used for fragmentation.

#### Database Search and Analysis

Data search was performed using MaxQuant (version 2.1.3.0) with an integrated Andromeda search engine, against the human Swissprot protein database (Downloaded on October 10^th^, 2022, containing 20,398 reviewed sequences). Digestion was defined as Trypsin/P and a maximum of 2 missed cleavages were allowed. Cysteine carbamidomethylation was set as a fixed modification. Protein N-terminal acetylation, methionine oxidation, and phosphorylation on serine, threonine, and tyrosine were set as variable modifications. Label-free quantification (LFQ) and the match-between-runs feature were enabled for protein quantification. A false discovery rate (FDR) of 1% was applied to both peptide spectrum matches (PSMs) and protein identification using a target-decoy approach. For total proteome measurements, intensity-based absolute quantification (iBAQ) was enabled.

Quantitative data filtering was conducted using the Perseus software (version 1.6.14.0). Proteins cross-matching to bovine contaminants were removed along with potential contaminants, reverse peptides, and proteins only identified by sites. LFQ intensities were log2-transformed. Proteins that were quantifiable in at least two out of three replicates were retained. Imputation was performed based on the normal distribution.

### Constitutive cell migration assessment

To assess the potential effect of CXCR4 cluster disruption on basal cell motility, Z-138 cells with a viability of > 90% were prepared at a concentration of 6 × 10^6^ cells/mL in standard growth (FBS-supplemented) media. These cells were treated with 1 μM of either IT1t, AMD3100, VUN401, VUN415 or vehicle. After an incubation period of 1 hour at 37°C, the cells were mixed 2:1 with ice cold BD Matrigel™. All plastics, including tips, eppindorf tubes and imaging slides were pre-chilled before use. Subsequently, 6 μL of the cell suspension was loaded into the central imaging chamber of an Ibidi µ- slide Chemotaxis (IbiTreat surface modification), according to manufacturer’s instruction. The Matrigel was allowed to solidify at 37°C for 30 minutes before filling the reservoirs flanking each chamber with media containing equal concentration of the compound.

Time-lapse video microscopy was conducted by capturing an image with a 10x phase contrast objective, every 5 minutes for 4 hours using a Nikon Ti2 microscope equipped with (37°C) temperature and (5%) CO2 control. Image analysis was performed using the open-source image processing software ImageJ2, version 2.14.0/1.54f. The manual tracking plugin was employed to analyze the trajectories of cells exhibiting high basal motility. Per condition 5 different cells were included in the conducted analysis. The Ibidi Chemotaxis and Migration Tool ImageJ plugin was utilized to generate Rose plots and extract average trajectory information.

### Resazurin assays for venetoclax sensitization

A total of 3 × 10^4^ Z-138, Jeko-1 and Maver-1 cells with a viability of > 90% were seeded in serum-free IMDM in a black 96-well plate (Greiner Bio-One). For primary cultures, 3 × 10^4^ PBMCs of CLL patients were thawed and seeded in IMDM supplemented with 10% FBS in a black 96-well plate. After 1 h, cells were treated with increasing concentrations of venetoclax in the absence or presence of 10 μM IT1t, AMD070, AMD3100, TG-0054 or VUN401. After 48 h incubation, 44 μM resazurin was added to the culture medium. After 1 h incubation, fluorescence cytotoxicity read-out was performed using a CLARIOstar plate reader at 540/30 nm excitation and 590/30 nm emission.

### FACS viability assays for venetoclax sensitization

A total of 3 × 10^4^ Z-138, Jeko-1 and Maver-1 cells with a viability of > 90% were seeded in serum-free IMDM in a transparent flat 96-well plate (Greiner Bio-One). After 1 h, cells were treated with increasing concentrations of venetoclax in the absence or presence of 10 μM IT1t, AMD070, AMD3100, TG-0054

or VUN401 and 20 μM pan-caspase inhibitor qVD-OPH. After 48 h incubation, 100 nM MitoTracker Orange (ThermoFisher Scientific, M7510) and 20 nM Topro-3 (ThermoFisher Scientific, T3605) were added according to the manufacturer’s guidelines. Well contents were transferred to polypropylene 5 mL tubes (Falcon) and analyzed using an Attune NxT Flow Cytometer.

Synergy assessment was done using the Bliss independence model, where ΔBliss scores for two compounds were calculated according to the following formulas:

(1) Δ*Bliss* = *E_Expected_ − E_Observed_*

(2) 
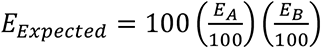

The Bliss independence model was used to assess whether the combined effect of compounds A and B is higher than the expected effect (E_Expected_) based on the relative individual effects (E_A_ and E_B_).

### Spheroid assays

PBMCs of CLL patients were thawed, plated in ultra-low attachment plates and centrifuged for 10 min at 1000 rpm and subsequently incubated for 24 h to allow spheroid formation. Three-dimensional (3D) cultures were cultured in IMDM supplemented with 10% FBS and 1% P/S and were stimulated and treated as indicated. Culture plates were placed in an IncuCyte live-cell imager (Essen Biosciences) in an incubator at 37⁰C and 5% CO2. Scans were taken every 5 h using the single spheroid assay for live-cell analysis application and 4x magnification. Spheroid area was quantified using IncuCyte software as a proxy for spheroid growth. Corresponding step-by-step protocols were previously described (*41*).

After culture, spheroids were resuspended and disintegrated to ensure proper antibody staining. Cells were incubated with monoclonal antibodies for surface staining for 30 min at 4⁰C. Cells were stained with antibodies against CD4 (AF700-labeled OK-T4, 56-0048-82, eBioscience), CD8 (PE- Cy7-labeled RPA-T8, 25-0088-42, eBioscience), CD19 (APC-labeled HIB19, 555415, BD Biosciences) and CD5 (PerCP-eF710-labeled UCHT2, 46-0059-42, eBioscience) for gaiting and with anti- CD25 (PE-conjugated clone M-A251, 555432, BD Biosciences) and Fixable Viability Dye eFluor™ 780 (ThermoFisher, 65-0865-14) to measure T cell activation and viability. Samples were measured on a Canto II flow cytometer (BD Biosciences). Samples were analyzed using FlowJo software.

### Data analysis

All graphs and bar plots were visualized, and statistical analyses were performed using Prism version 10.0 (GraphPad) unless indicated otherwise. Curves were fitted using least squares nonlinear regressions, assuming a sigmoidal fit (for concentration-response curves). The significance of differences was determined as indicated in the figure legends. Schematics for assay formats were generated using Biorender.com

### Data Availability

The phosphoproteomics data have been deposited to the ProteomeXchange Consortium through the PRIDE partner repository with the dataset identifier PXD053673.

## Acknowledgements

This work was funded by the Netherlands Organization for Scientific Research (NWO) grant ENPPS.TA.019.003 MAGNETIC for S.M, N.D.B. and Z.M., M.Sp., W.W., S.H.T., A.P.K. and R.H., the ZonMw Veni grant 09150162010212 for C.C., the ZonMw grant 91217002 for D.J., the European Union H2020-MSCA grant 641833-ONCORNET for M.Si., M.J.S. and R.H., H2020-MSCA grant 860229-ONCORNET 2.0 for S.M.A., Z.W. and C.V.P.A., M.L., M.Si., M.J.S. and R.H. We greatly acknowledge the Microscopy & Cytometry core facility (MCCF) of the Amsterdam University Medical Center for imaging and cytometry support. We greatly appreciate Prof. Dr. Lukas Kapitein for helping out with the dSTORM microscopy experiments.

