## Supplemental Information for "Disruption of basal CXCR4 oligomers impairs oncogenic properties in lymphoid neoplasms"

**Supplementary Information**

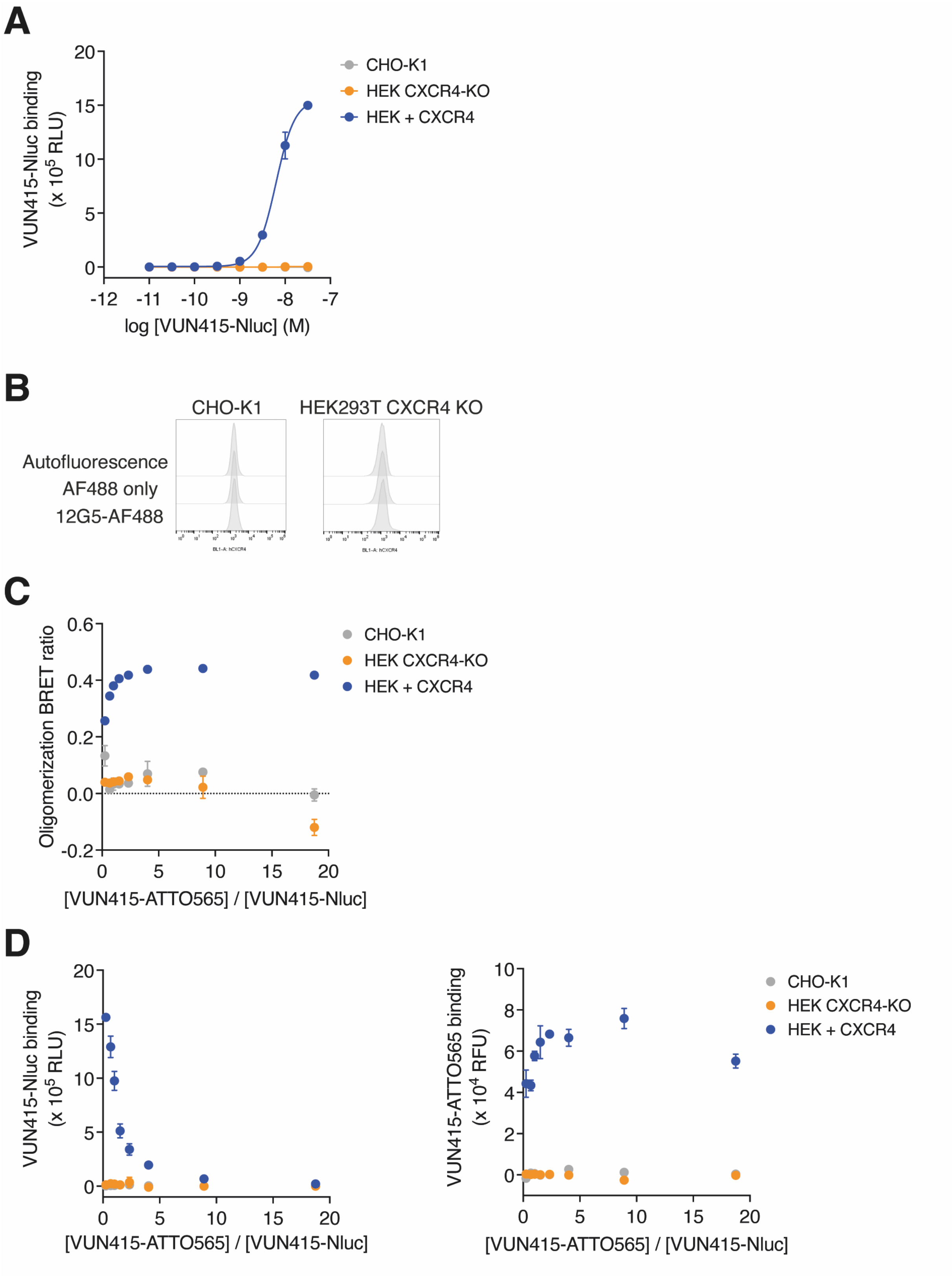

**Figure S1. Controls for nanobody-based BRET to detect oligomers of transfected CXCR4. A** VUN415-NanoLuc binding curves in CHO-K1, HEK293T CXCR4 CRISPR Cas9 KO and CXCR4-overexpressing HEK293T cells. **B** CXCR4 surface expression levels in CHO-K1 and HEK293T CXCR4 KO cell lines detected by flow cytometry with CXCR4 antibody 12G5. In the same experiment, autofluorescence was determined and cells were treated with secondary antibodies alone as control conditions. **C** Nanobody-based saturation setup measurement of CXCR4 oligomerization in CHO-K1, HEK293T CXCR4 CRISPR Cas9 KO and CXCR4-overexpressing HEK293T cells. 31.6 nM detection nanobodies with increasing ratios of VUN415-ATTO565 to VUN415-NanoLuc were used for oligomer detection. **D** Binding curves of VUN415-NanoLuc and VUN415-ATTO565 in parallel of oligomerization detection experiment in CHO-K1, HEK293T CXCR4 CRISPR Cas9 KO and CXCR4-overexpressing HEK293T cells. Data are mean ± SD and are representative of three independent experiments, each performed in triplicate **(A, C, D)** or are representative histograms of three independent experiments **(B)**.

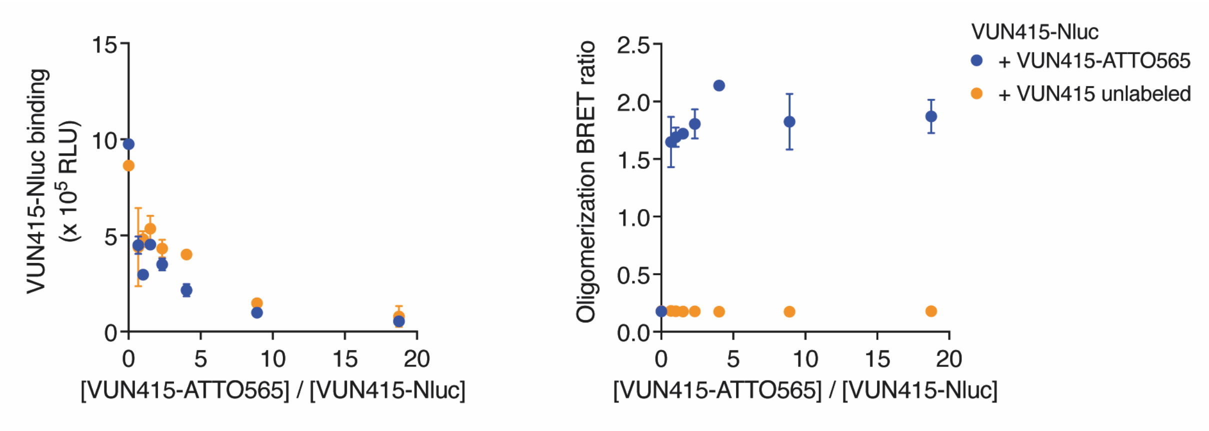


**Figure S2. No oligomerization BRET signal when using VUN415-NanoLuc with unlabeled VUN415 instead of VUN415-ATTO565.** VUN415-NanoLuc binding and nanobody-based saturation setup measurement of CXCR4 oligomerization in CXCR4-overexpressing HEK293T cells. 31.6 nM nanobodies with increasing ratios of VUN415(-ATTO565) to VUN415-NanoLuc were used for oligomer detection. Data are mean ± SD and are representative of three independent experiments, each performed in triplicate.

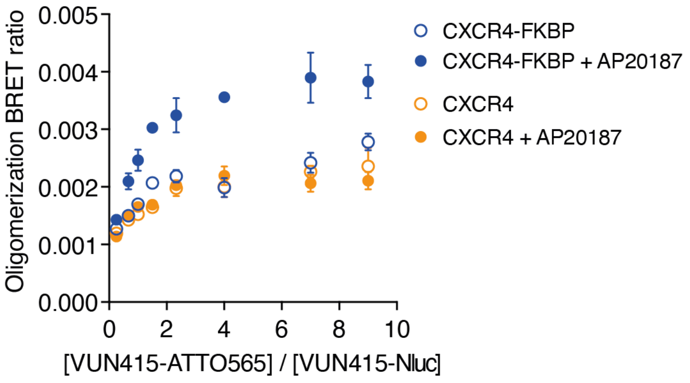


**Figure S3. The FKBP domain demonstrates the specificity of the nanobody-based method to detect forced dimerization at low expression levels.** Nanobody-based saturation setup measurement of CXCR4 oligomerization in CXCR4 and CXCR4-FKBP overexpressing HEK293T cells. Cells were stimulated with AP201827 (1 uM) to induce receptor dimerization of CXCR4-FKBP. 31.6 nM detection nanobodies with increasing ratios of VUN415-ATTO565 to VUN415-NanoLuc were used for oligomer detection. Data are mean ± SD and are representative of three independent experiments, each performed in triplicate

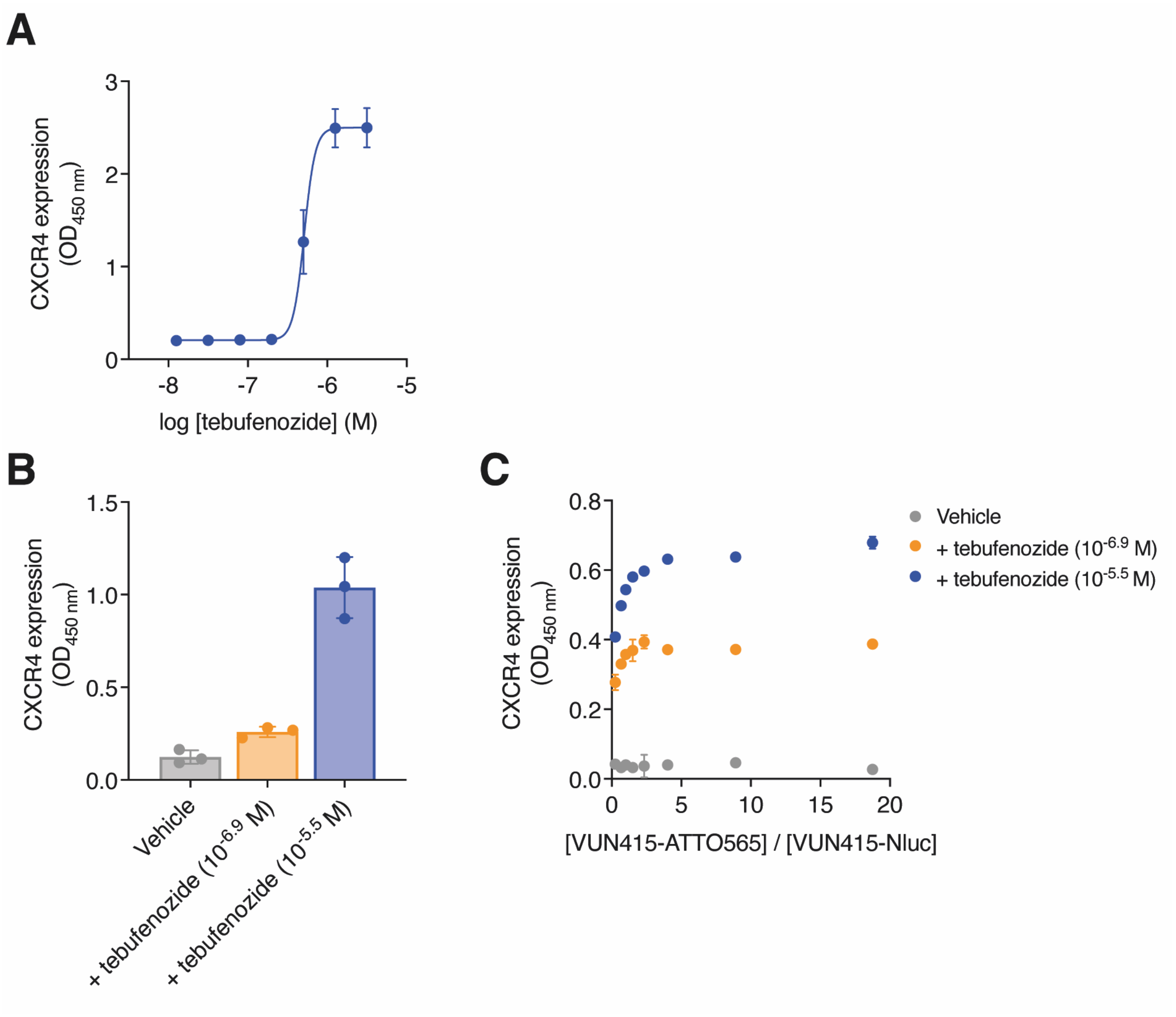


**Figure S4. Nanobody-based BRET method validates expression dependency for oligomerization of transfected CXCR4. A, B** ELISA-based measurement of receptor expression levels for ecdysone-inducible CXCR4 in HEK293T cells using equimolar **(A)** and saturation setup **(B)**. Stimulation with increasing concentrations tebufenozide to induce receptor expression. **C** Nanobody-based saturation setup measurement of ecdysone-inducible CXCR4 oligomerization in HEK293T cells. 31.6 nM detection nanobodies with increasing ratios of VUN415-ATTO565 to VUN415-NanoLuc were used for oligomer detection. Data are mean ± SD and are representative of three independent experiments, each performed in triplicate.

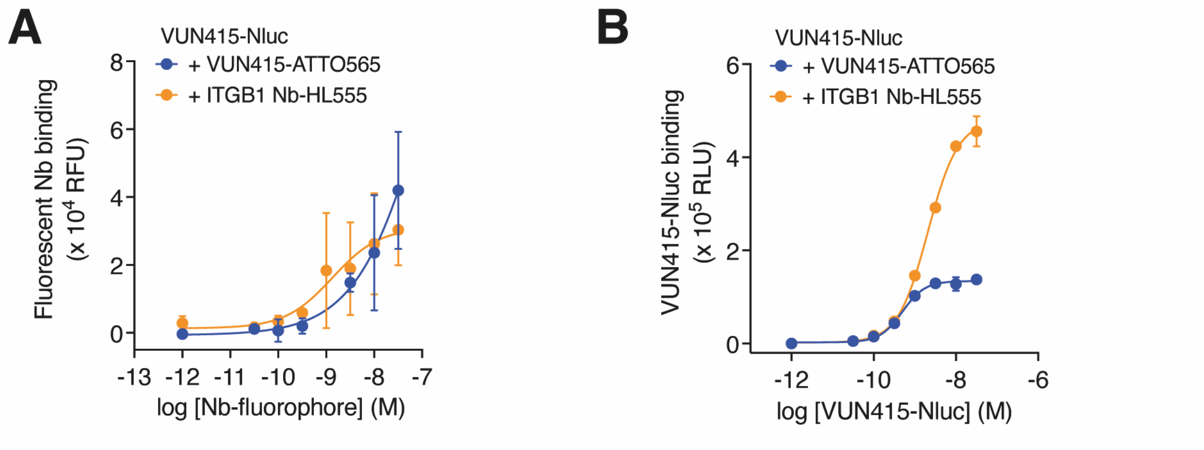


**Figure S5**. **Absence of oligomerization BRET signal is not due to lack of ITGB1-Nb-HL555 binding. A, B** Binding curves of increasing concentrations VUN415-ATTO565 and ITGB1-Nb-HL555 **(A)** or VUN415-NanoLuc **(B)** in parallel of oligomerization detection experiment in Namalwa cells. Data are mean ± SD and are representative of three independent experiments, each performed in triplicate.

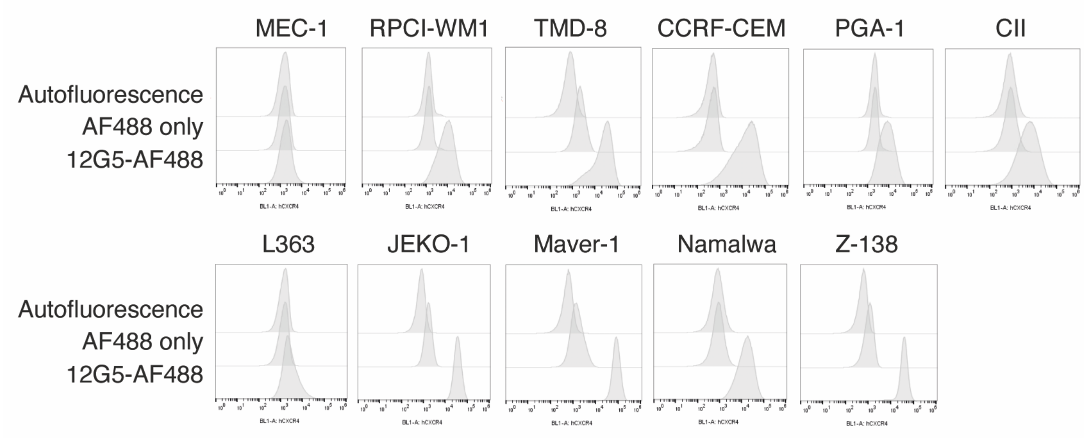


**Figure S6. CXCR4 expression levels for the complete panel of lymphoid cancer cell lines.** CXCR4 surface expression levels in the panel of lymphoid cancer cell lines detected by flow cytometry with CXCR4 antibody 12G5. In the same experiment, autofluorescence was determined and cells were treated with secondary antibodies alone as control conditions. Data are representative histograms of three independent experiments.

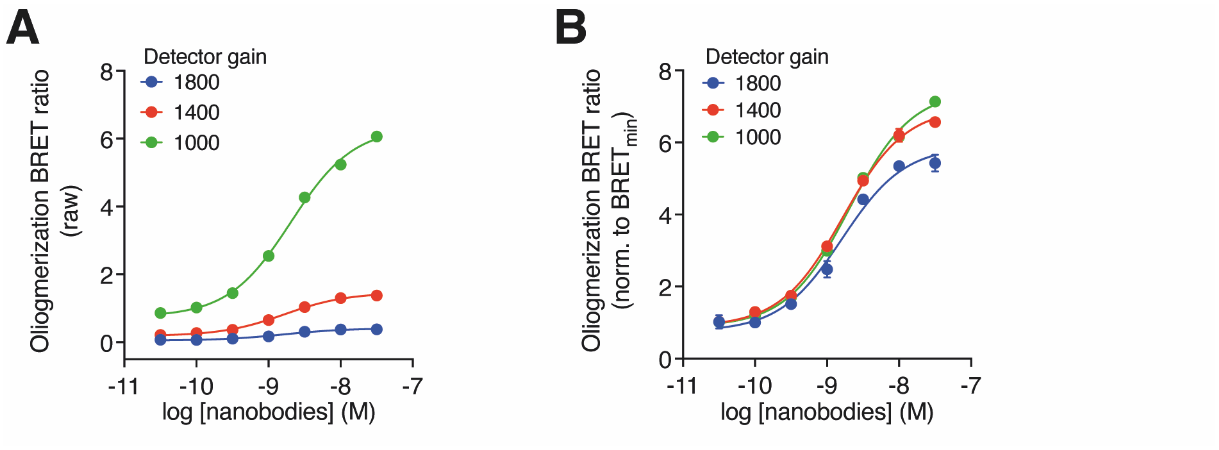


**Figure S7. Plate reader gain settings do not influence normalized oligomerization $BRET_{max}$ values.** Nanobody-based equimolar setup measurement of CXCR4 oligomerization in Z-138 cells. Increasing equimolar concentration of detection nanobodies VUN415-NanoLuc and VUN415-ATTO565 were used. Curves of raw **(A)** and $BRET_{min}$-normalized (**B)** BRET ratios for different gain settings are shown. Data are mean ± SD and are representative of three independent experiments, each performed in triplicate.

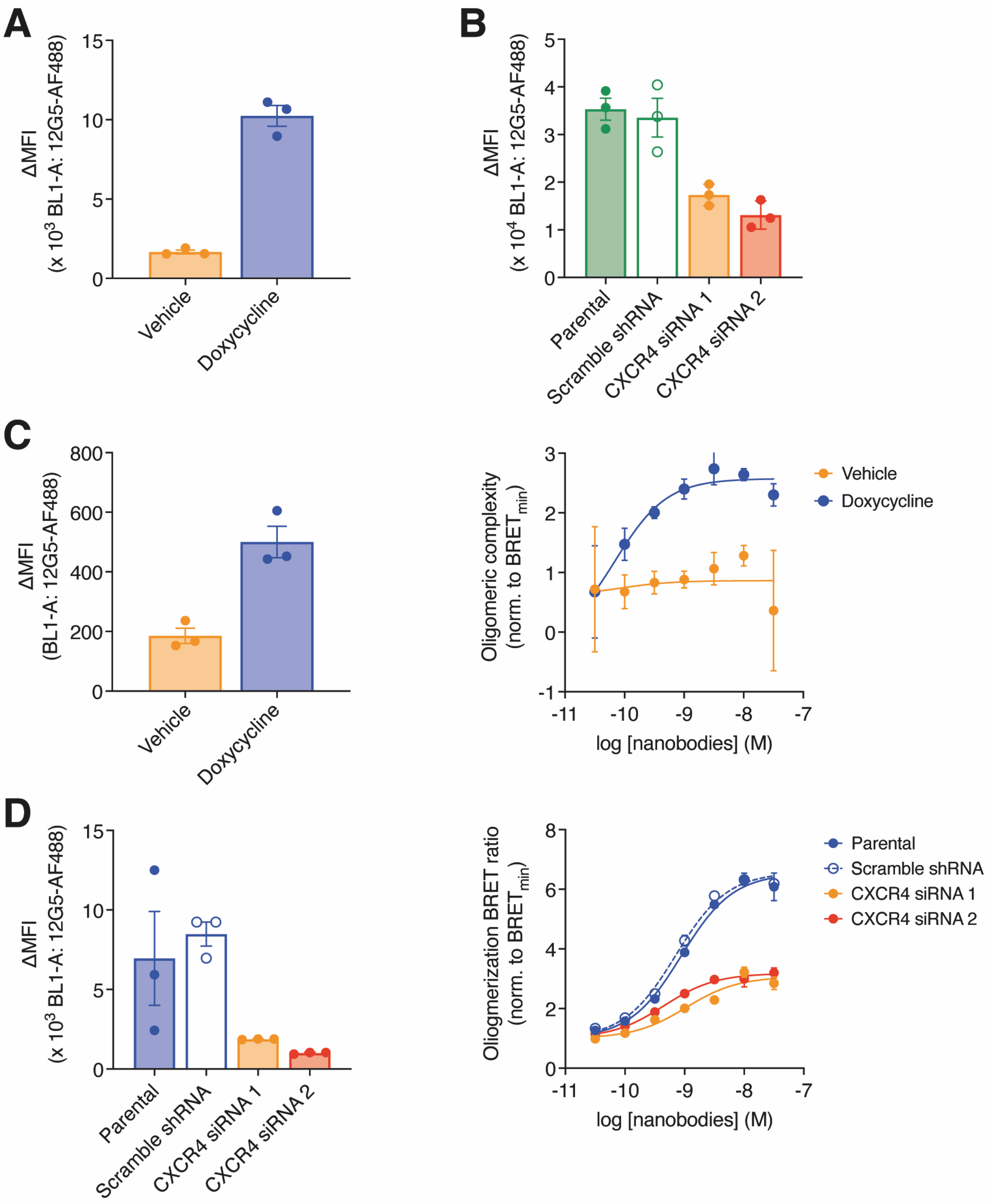


**Figure S8. Validation of knockdown and upregulation in genetic approaches to modulate CXCR4 oligomerization. A, B** CXCR4 surface expression levels detected by flow cytometry with CXCR4 antibody 12G5 in doxycycline stimulated RPCI-WM1 + iCXCR4 cells **(A)** or Z-138 ± scramble shRNA or CXCR4 siRNA-1/2 cells **(B)**. Delta MFI was calculated by subtracting geometric MFI from the secondary antibody-only condition. Data are the pooled mean ± SEM of three independent experiments, each performed in duplicate. **C, D** CXCR4 surface expression levels and nanobody-based BRET measurement of oligomeric complexity in doxycycline stimulated MEC-1 iCXCR4 cells **(C)** or Namalwa ± scramble shRNA or CXCR4 siRNA-1/2 cells **(D)**. Increasing equimolar concentrations of VUN415-NanoLuc and VUN415-ATTO565 were used for oligomer detection. Data are the mean ± SD and are representative of three independent experiments, each performed in triplicate.

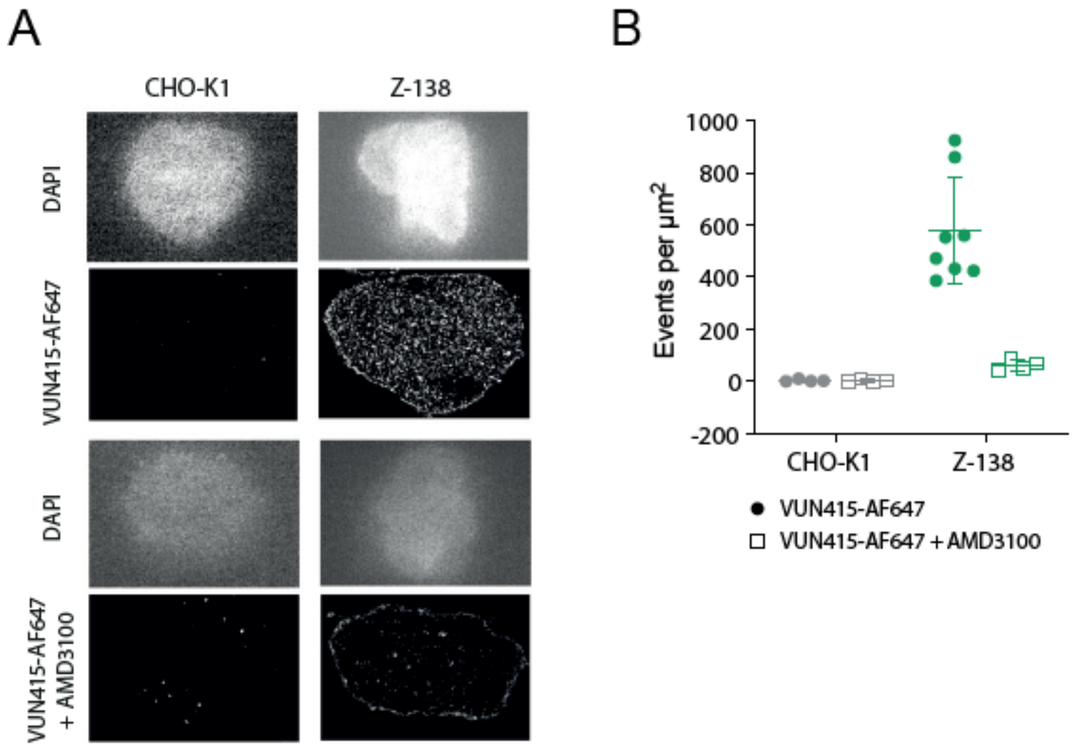


**Figure S9. Additional dSTORM data. A** Reconstructed dSTORM images of VUN415-Alexa647 stains in the presence or absence of excess AMD3100 and corresponding DAPI stains of CHO-K1 and Z-138 cells. **B** CXCR4 density assessment on CHO-K1 and Z-138 cells. Pooled mean fractions ± SEM are of four or eight analyzed areas, from two independent experiments per cell line.

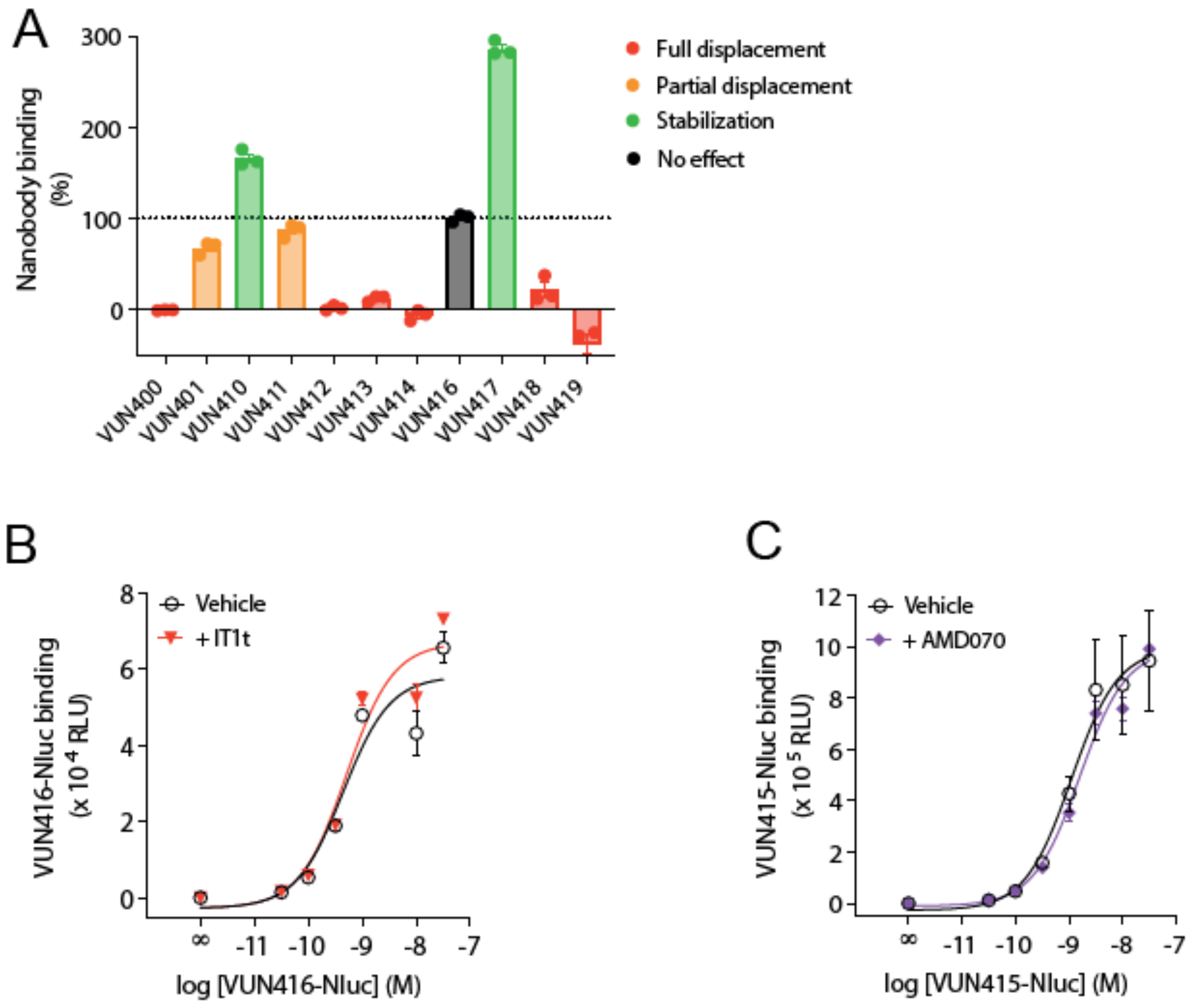


**Figure S10. Screen identifies VUN416 as non-IT1t-competitive nanobody and VUN415 as non-AMD070 competitive nanobody.** BRET-based measurement of indicated nanobody-ATTO565 (1 nM) binding in presence of IT1t (10 µM) using membrane extracts from NanoLuc-CXCR4-expressing HEK293T cells. Dotted line indicates the fluorescent ligand only condition. Data, normalized to the fluorescent ligand only condition, are the pooled mean ± SEM of three independent experiments, each performed in duplicate. **B** Binding curves of increasing concentrations VUN416-NanoLuc in the absence or presence IT1t (10 µM), in parallel with the oligomerization detection experiments in Z-138 cells. Data are mean ± SD and are representative of three independent experiments, each performed in triplicate. **C** Binding of increasing concentrations of VUN415-NanoLuc in the absence or presence of AMD070 (10 µM), in parallel of the oligomerization detection experiment in Z-138 cells. Data, normalized to the buffer-only condition, are the mean ± SD and are representative of three independent experiments, each performed in triplicate.

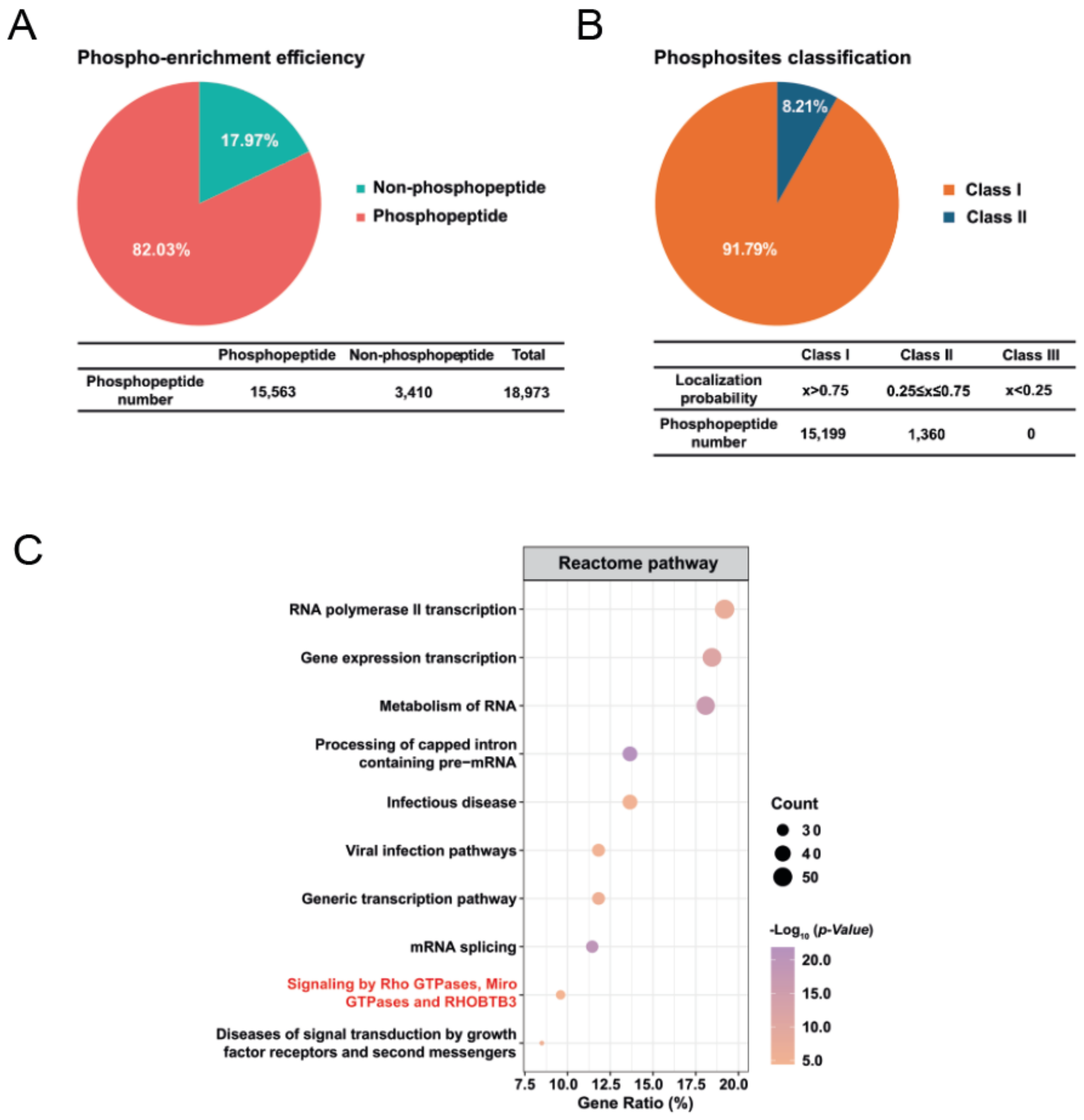


**Figure S11. Quality control of phosphoproteomics samples. A and B** Evaluation of phospho-enrichment efficiency (A) and phospho-site classification (B). **C** Biological pathways modulated by CXCR4 cluster disruption by IT1t. Evaluation of the pathways modulated by cluster disruptors IT1t following stimulation of 60 minutes, displayed as Gene Ontology plot.

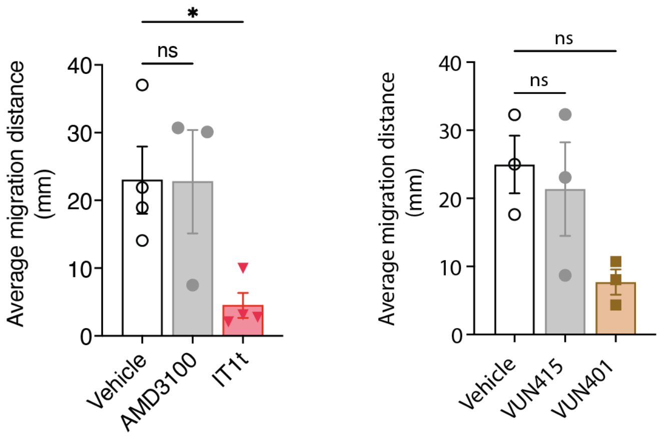


**Figure S12. CXCR4 monomerizing ligands impair the basal migration of an MCL cell line.** The average traveled distance of MCL Z-138 cells after four hours following treatment with 1 μM of AMD3100, IT1t, VUN415 or VUN401. Data are pooled mean ± SEM of at least three independent experiments. * $P < 0.05$ compared to vehicle, according to one-way ANOVA followed by Dunnet's post hoc test.

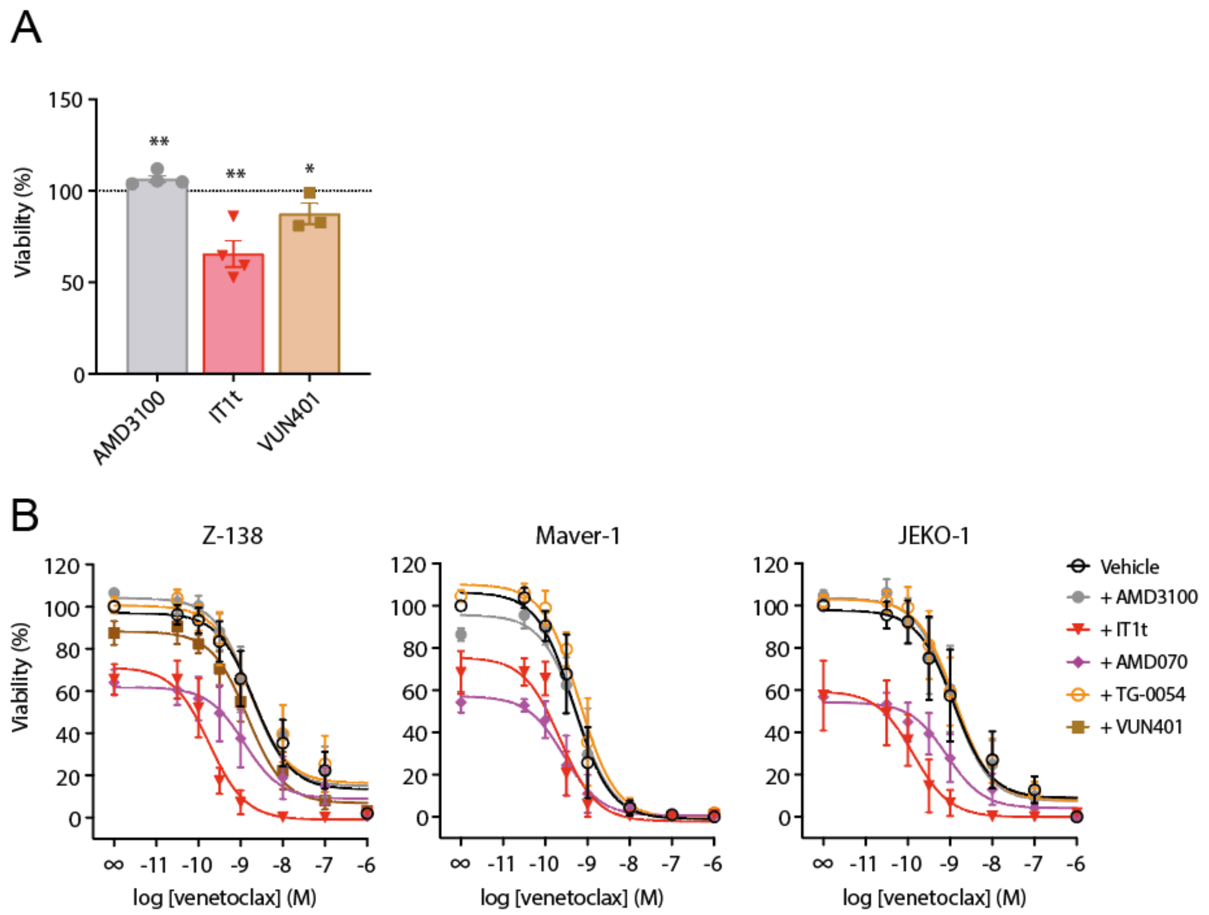


**Figure S13. CXCR4 monomerizing ligands inhibit viability of Z138, Maver-1 and JEKO-1 cells. A** FACS-based measurement of cell death in MCL Z-138 cells after 48 h treatment in absence (vehicle) or presence of indicated ligands (10 µM). Data are pooled mean ± SEM of at least three independent experiments. **B** FACS-based measurement of cell death in indicated lymphoid cancer cell lines after 48 h treatment with venetoclax in the absence (vehicle) or presence of indicated ligands (10 µM). Data are pooled mean ± SEM of at least three independent experiments. Data, normalized to the 'no venetoclax' condition, are pooled mean ± SEM of at least three independent experiments, each performed in triplicate.

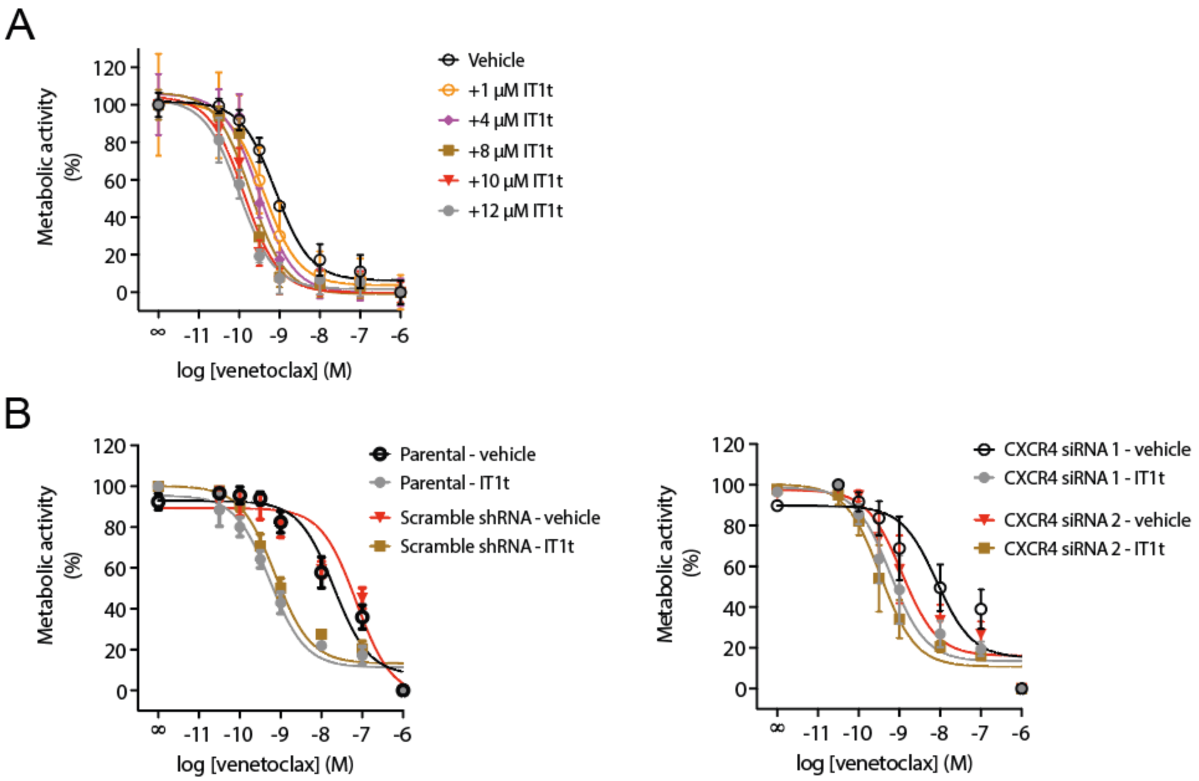


**Figure S14. Sensitization of MCL cell line Z-138 to venetoclax depends on IT1t dose and and high CXCR4 expression. A** Resazurin-based measurement of metabolic activity in Z-138 cells after 48 h treatment with increasing concentrations of venetoclax in absence (vehicle) or presence of five increasing concentrations IT1t. Data, normalized to the 'no venetoclax, vehicle' condition, are pooled mean ± SEM of at least three independent experiments, each performed in triplicate. **B** Resazurin-based measurement of metabolic activity in Z-138 upon CXCR4 knock down (right) or negative controls (left) and 48 h treatment with a concentration range of venetoclax in absence (vehicle) or presence of IT1t (10 µM). Data, normalized to the 'no venetoclax' condition, are pooled mean ± SEM of at least three independent experiments, each performed in triplicate. * $P < 0.05$, ** $P < 0.01$, compared to vehicle, according to unpaired t-test.

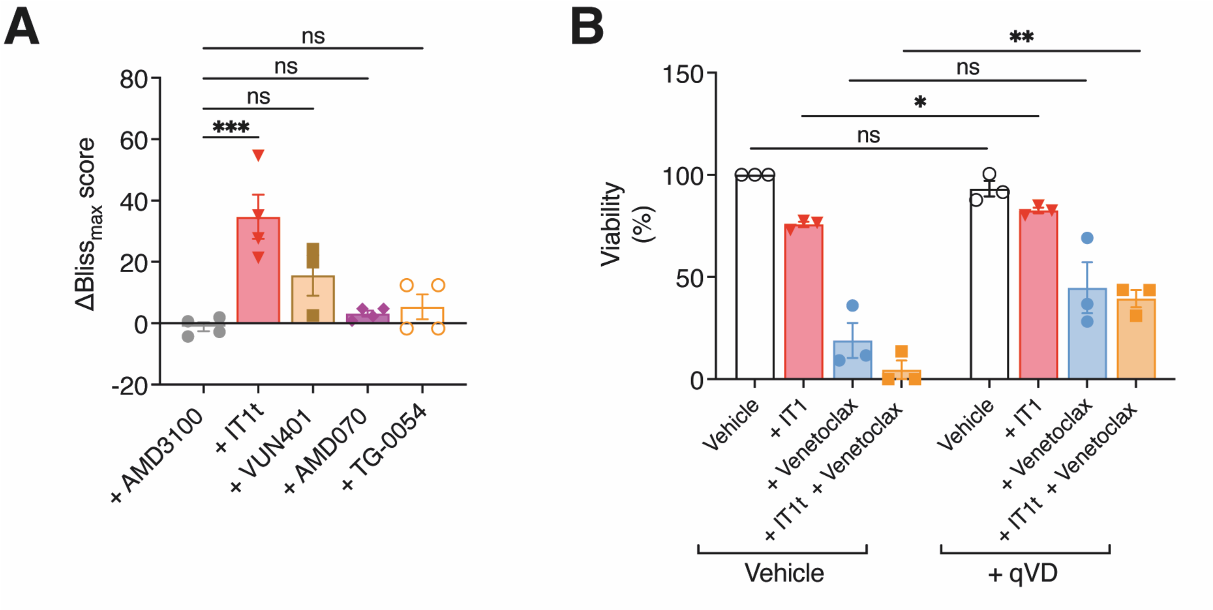


**Figure S15. CXCR4-monomerizing ligands sensitize multiple MCL cell lines to venetoclax-induced apoptosis in a synergistic manner. A** ΔBliss score assessment on the FACS-based cell death measurements in Z-138 after 48 h treatment with venetoclax in the absence (vehicle) or presence of indicated ligands (10 µM). Maximal ΔBliss scores ($\Delta Bliss_{max}$) at variable venetoclax concentrations are displayed (100 nM venetoclax for VUN401, 1 nM venetoclax for AMD070 and AMD3100, 316 pM venetoclax for IT1t and 100 pM venetoclax for TG-0054). Data are pooled mean ± SEM of at least three independent experiments. **B** FACS-based measurement of cell death in Z-138 after 48 h treatment with venetoclax (3 nM), IT1t (10 µM) or combined in the absence or presence of the pan-caspase inhibitor qVD-OPH (20 µM). Data are pooled mean ± SEM of at least three independent experiments.

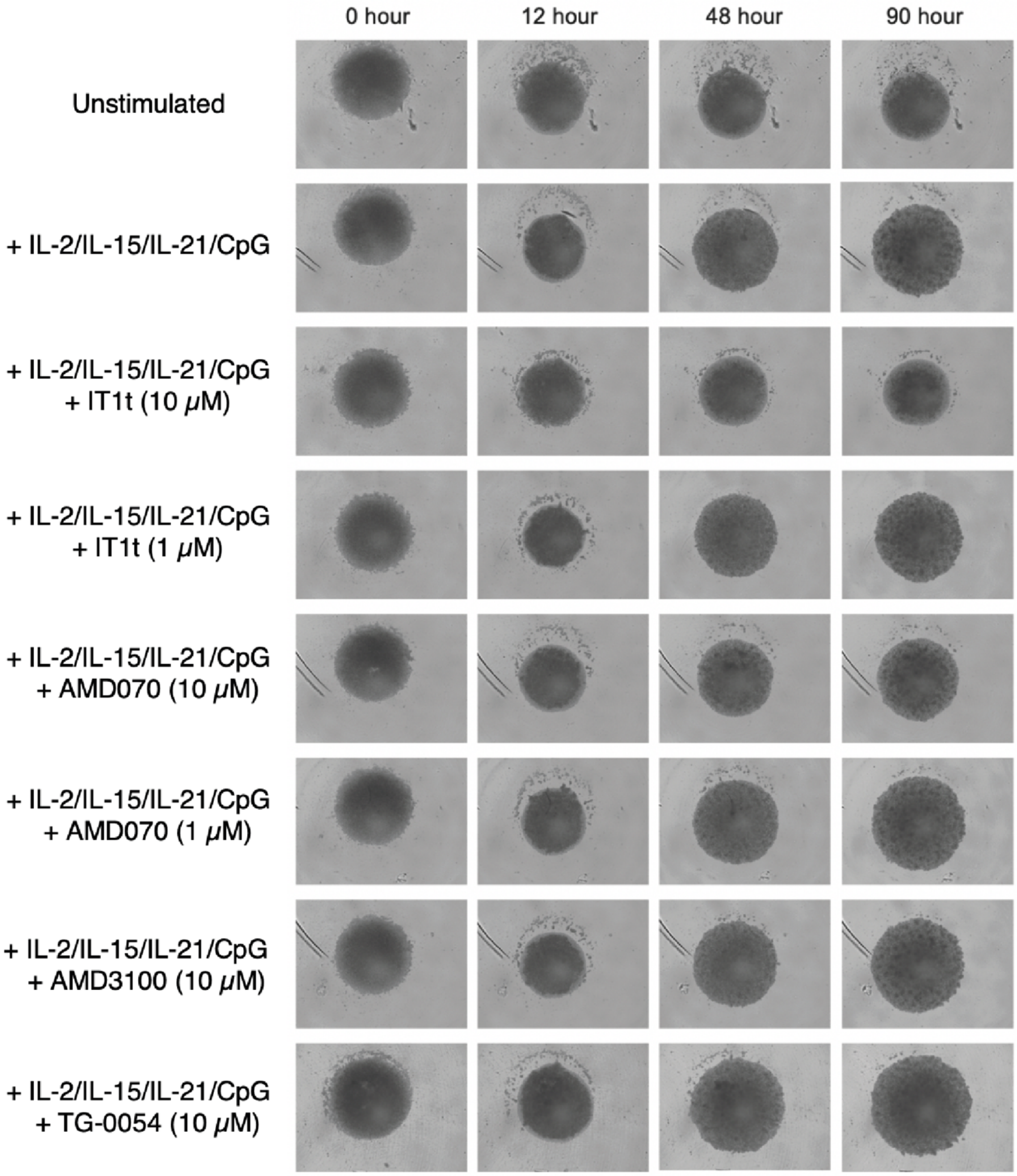


**Figure S16. CXCR4-monomerizing ligands selectively inhibit growth in a CLL patient-derived spheroid model.** Effects of indicated CXCR4 antagonists on IL-2/IL-15/IL-21/CpG cocktail-induced growth curve in CLL patient-derived spheroid model (*41*). Representative images of a culture derived from a single patient are shown.

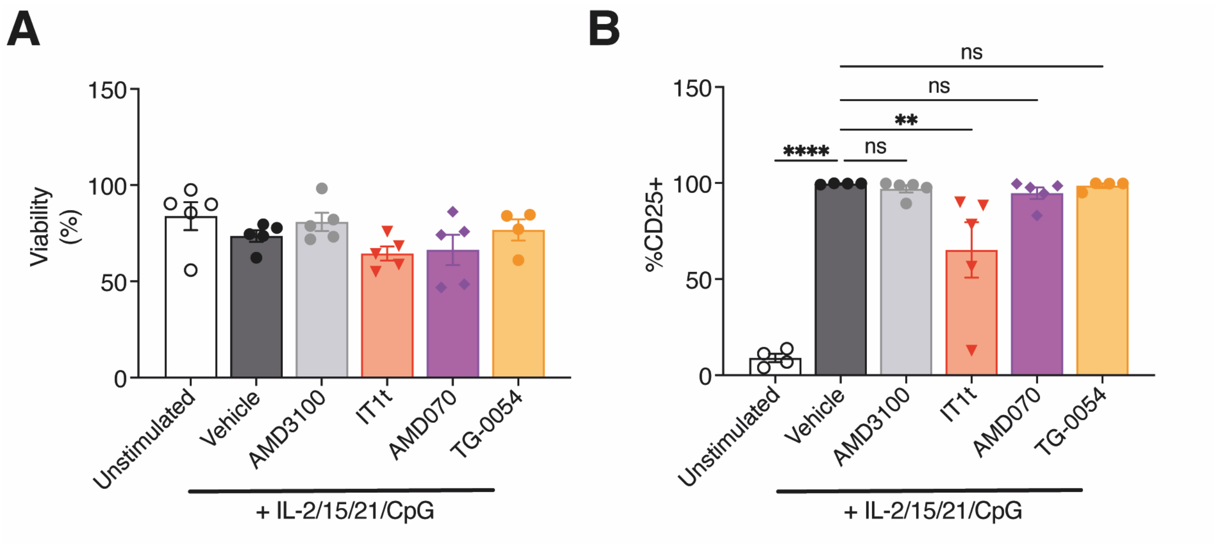


**Figure S17. Spheroid growth inhibition is not due to toxicity of compounds. A, B** Effects of indicated CXCR4 antagonists (10 µM) on cell viability (**A**) and expression of activation marker CD25 **(B)** in primary CLL spheroid model after 90 hours of treatment. Data are mean ± SEM of cultures from four (TG-0054) or five individual patients. No significant difference compared to vehicle, according to one-way ANOVAs followed by Dunnet's post hoc test **(A)**. ** $P < 0.01$, **** $P < 0.0001$, compared to vehicle according to one-way ANOVAs followed by Dunnet's post hoc test **(B)**.

**Table S1. Screen for CXCR4-targeting oligomer detection nanobody with high receptor binding affinity and neutral properties.** Data, normalized to the fluorescent ligand only (for CXCL12-AF647 displacement) or the vehicle condition (for modulation CXCR4 oligomerization and inhibition of basal $G\alpha_{i/o}$ activation), are pooled mean ± SEM of three independent experiments, each performed in duplicate. Significant differences (P < 0.05) compared to vehicle condition are indicated in bold, according to one-sample t-test.

| | **CXCL12-AZ647 displacement** | | **Modulation CXCR4 oligomerization** | | **Inhibition basal $G\alpha_{i/o}$ activation** | |
|---|---|---|---|---|---|---|
| **Nanobody** | Displacement, % (mean ± SEM) | $pIC_{50}$ (mean ± SEM) | $E_{max}$ (mean ± SEM) | P | $E_{max}$ (mean ± SEM) | P |
| **VUN400** | 99 ± 5 | 8.1 ± 0.11 | 1.01 ± 0.016 | 0.652 | **0.90 ± 0.012** | **0.014** |
| **VUN401** | 106 ± 9 | 7.1 ± 0.16 | **0.87 ± 0.011** | **0.008** | 1.04 ± 0.013 | 0.088 |
| **VUN410** | 92 ± 3 | 8.0 ± 0.08 | 0.99 ± 0.013 | 0.514 | 1.00 ± 0.009 | >0.999 |
| **VUN411** | 100 ± 4 | 8.1 ± 0.09 | **0.94 ± 0.003** | **0.002** | 0.99 ± 0.004 | 0.163 |
| **VUN412** | 101 ± 3 | 8.3 ± 0.09 | 1.00 ± 0.015 | 0.999 | 0.98 ± 0.011 | 0.225 |
| **VUN413** | 103 ± 7 | 7.8 ± 0.16 | 1.02 ± 0.018 | 0.380 | **0.93 ± 0.009** | **0.15** |
| **VUN414** | 100 ± 5 | 8.0 ± 0.13 | 0.98 ± 0.007 | 0.117 | **0.93 ± 0.001** | **0.0003** |
| **VUN415** | 102 ± 5 | 8.7 ± 0.13 | 0.99 ± 0.004 | 0.163 | 1.01 ± 0.003 | 0.074 |
| **VUN416** | 105 ± 5 | 8.8 ± 0.12 | 0.98 ± 0.007 | 0.117 | **0.97 ± 0.002** | **0.042** |
| **VUN417** | 107 ± 6 | 7.5 ± 0.13 | 1.02 ± 0.010 | 0.194 | 0.99 ± 0.004 | 0.102 |
| **VUN418** | 108 ± 6 | 7.4 ± 0.13 | **0.90 ± 0.008** | **0.002** | 1.04 ± 0.010 | 0.165 |
| **VUN419** | 112 ± 6 | 8.4 ± 0.17 | **0.90 ± 0.019** | **0.012** | 1.01 ± 0.016 | 0.559 |

**Table S2. Effects CXCR4 ligands on venetoclax potency in different MCL and CLL cell lines in a resazurin readout.** Data are pooled mean ± SEM of at least three independent experiments, each performed in triplicate. Significant differences (P < 0.05) compared to vehicle condition are indicated in bold, according to a one-way ANOVA followed by Dunnet's post hoc test. * P < 0.05, ** P < 0.01, **** P < 0.0001.

| | **Z-138** | | **Maver-1** | | **Jeko-1** | | **PGA-1** | |
|---|---|---|---|---|---|---|---|---|
| **Treatment** | $pIC_{50}$ | P | $pIC_{50}$ | P | $pIC_{50}$ | P | $pIC_{50}$ | P |
| **Vehicle** | 9.3 ± 0.1 | N.A. | 9.6 ± 0.1 | N.A. | 8.2 ± 0.1 | N.A. | 6.6 ± 0.7 | N.A. |
| **AMD3100** | 9.3 ± 0.1 | 0.99 | 9.6 ± 0.1 | 0.93 | 8.1 ± 0.3 | 0.95 | 6.3 ± 1.3 | 0.95 |
| **IT1t** | **10.1 ± 0.1** | **** | **10.0 ± 0.1** | ** | **8.9 ± 0.1** | * | 9.7 ± 0.4 | 0.08 |
| **VUN401** | **9.7 ± 0.2** | ** | N.D. | N.D. | N.D. | N.D. | N.D. | N.D. |

**Table S3. Effects CXCR4 ligands on venetoclax potency in different MCL and CLL cell lines in a FACS viability readout.** Data are pooled mean ± SEM of at least three independent experiments, each performed in triplicate. Significant differences ($P < 0.05$) compared to vehicle condition are indicated in bold, according to a one-way ANOVA followed by Dunnet's post hoc test. * $P < 0.05$, ** $P < 0.01$.

| | **Z-138** | | **Maver-1** | | **Jeko-1** | |
|---|---|---|---|---|---|---|
| **Treatment** | $p\mathrm{IC}_{50}$ | P | $p\mathrm{IC}_{50}$ | P | $p\mathrm{IC}_{50}$ | P |
| **Vehicle** | 8.5 ± 0.6 | N.A. | 9.3 ± 0.3 | N.A. | 8.8 ± 0.6 | N.A. |
| **IT1t** | **9.7 ± 0.4** | * | 9.6 ± 0.3 | 0.1 | **9.8 ± 0.5** | * |
| **AMD070** | **8.8 ± 0.7** | ** | 9.5 ± 0.5 | 0.2 | 9.0 ± 0.7 | 0.3 |
| **TG-0054** | 8.3 ± 1 | 0.3 | 9.2 ± 0.3 | 0.1 | 8.9 ± 0.4 | 0.8 |
| **AMD3100** | 8.3 ± 0.9 | 0.3 | 9.2 ± 0.4 | 0.3 | 8.9 ± 0.5 | 0.4 |
| **VUN401** | 8.8 ± 0.3 | 0.1 | N.D. | N.D. | N.D. | N.D. |

**Table S4. Patient information table.** Information about the used CLL and MCL patient samples in this study.

| **Malignancy** | **Sample** | **Sample date** | **Age** | **Gender** | **IGHV** | **FISH/mutations** | **Treatment** |
|---|---|---|---|---|---|---|---|
| CLL | 1961 | 16-06-2017 | 84 | M | Mutated | Trisomy 12 | Prednisone |
| | 2148 | 19-03-2018 | 81 | M | Mutated | del(13q14), TP53 mutation (VAF 1%) | none |
| | 1576 | 11-12-2014 | 88 | F | Mutated | Not performed | none |
| | 2094 | 23-01-2018 | 64 | F | Unmutated | trisomy 12, del(4p15.2-p15.1), TP53 WT | none |
| | 2748 | 18-11-2019 | 65 | F | Unmutated | trisomy 12, del(4p15.2-p15.1), TP53 WT | none |
| | 2747 | 14-11-2019 | 63 | M | Unmutated | Absence del11q, absence del17p, TP53 WT | none |
| | 664 | 12-06-2008 | 85 | M | Mutated | del(13q14) 58%, del(14q32)(IGH) 9% | none |
| | 50 | 15-10-2004 | 64 | F | Mutated | Not performed | none |
| MCL | 2257 | 14-06-2018 | 57 | M | ND | T(11;14) 80%, del(13q14) (loss of DLEU2/MIR15A/MIR16-1 and RB1 locus), del(17p13.1) (TP53), del(2q33.2-q34), dup(5q33.2-q34), dup(8p23.1-p22), del(9p24.1-p21.3), del(10p15.3-p13), 13q12.11-q13.3), chromotrypsis can chr. 14, dup(16q21-q22), dup(22q12.3-q13.2) | 2015: Acerta Btk inh. 20-06-2018: R Benda. |
| | No sample number available, only date | 19-07-2016 | 68 | M | ND | t (11;14) in 78% cells, complex karyotype with gain MYC. | Rasburicase & levocetirizine. Patient left ACERTA trial (acelabrutinib) just before sample. Had 6 cycles of acalabrutinib with comple remission. |
| | No sample number available, only date | 11-07-2014 | 79 | F | ND | T(11;14)(q13;q32) 20%, cycline d1 translocatie | 1st cycle Benda 17-07-2014. |
